# Glycine receptor α3K governs mobility and conductance of L/K splice variant heteropentamers

**DOI:** 10.1101/2021.02.18.431627

**Authors:** Veerle Lemmens, Bart Thevelein, Svenja Kankowski, Hideaki Mizuno, Jochen C. Meier, Susana Rocha, Bert Brône, Jelle Hendrix

**Author notes:** These authors contributed equally (Veerle Lemmens, Bart Thevelein). Correspondence, Hasselt University, Agoralaan C (BIOMED), B-3590 Diepenbeek.

## Abstract

Glycine receptors (GlyRs) are ligand-gated pentameric chloride channels in the central nervous system. GlyR-α3 is a possible target for chronic pain treatment and temporal lobe epilepsy. Alternative splicing into K or L variants determines the subcellular fate and function of GlyR-α3, yet it remains to be shown whether its different splice variants can functionally co-assemble, and what the properties of such heteropentamers would be. Here, we subjected GlyR-α3 to a combined fluorescence microscopy and electrophysiology analysis. We employ masked Pearson’s and dual-color spatiotemporal correlation analysis to prove that GlyR-α3 splice variants heteropentamerize, adopting the mobility of the K variant. Fluorescence-based single-subunit counting experiments revealed a variable and concentration ratio dependent hetero-stoichiometry. Via single-channel on-cell patch clamp we show heteropentameric conductances resemble those of the α3K splice variant. Our data are compatible with a model where α3 heteropentamerization fine-tunes mobility and activity of GlyR α3 channels, which is important to understand and tackle α3 related diseases.

**Summary:** The glycine receptor α3 is key to the central nervous system’s physiology and involved in chronic pain and epilepsy. In this paper, Lemmens et al. reveal and functionally characterize α3 splice variant heteropentamerization via advanced single-molecule fluorescence image analysis.

**Declarations:** *Funding:* We acknowledge the UHasselt Advanced Optical Microscopy Centre (AOMC). Prof. Em. Marcel Ameloot, the Research Foundation Flanders (FWO, project G0H3716N) and the province of Limburg (Belgium) (tUL Impuls II) are acknowledged for funding the microscopy hardware. V. Lemmens is grateful for a doctoral scholarship from the UHasselt (17DOC11BOF) and KU Leuven (C14/16/053) Special Research Funds (BOF).

*Conflicts of interest / competing interests:* No conflicts of interest apply.

*Ethics approval:* Not applicable

*Availability of data and material:* All data and material are available upon request.

*Code availability:* Fluctuation imaging and co-localization analyses were performed in the software package PAM [71]. The software is available as source code, requiring MATLAB to run, or as pre-compiled standalone distributions for Windows or MacOS at http://www.cup.uni-muenchen.de/pc/lamb/software/pam.html or hosted in Git repositories under http://www.gitlab.com/PAM-PIE/PAM and http://www.gitlab.com/PAM-PIE/PAMcompiled. A detailed user manual is available at http://pam.readthedocs.io.

*Author contributions:* Conceptualization Meier J.C., Brône B. and Hendrix J.; Investigation and formal analysis Lemmens V. and Thevelein B.; Software development Hendrix J.; Writing the original draft Lemmens V., Thevelein B and Hendrix, J.; Review and editing by all authors.

## Introduction

Neuronal communication in the central nervous system (CNS) is fine-tuned via ionotropic channel proteins such as glycine receptors (GlyRs). Belonging to the family of pentameric ligand-gated ion channels (pLGICs), GlyRs help regulate motor coordination and sensory signal processing [1, 2]. In humans, GlyRs are expressed as one of three α isoforms (α1-3) that heteropentamerize with the β isoform if the latter is present. In this paper we focus on the α3 isoform of GlyR, which in the human body is found in the spinal cord’s dorsal horn, the brain stem and the hippocampus[3]. In addition, high RNA levels were also found in the cerebral cortex, the amygdala and in the pituitary gland [4]. It is involved in temporal lobe epilepsy (TLE) [5–8] and, due to its crucial involvement in inflammatory pain perception, it is a major target for chronic pain treatment [9]. Because of its specific localization in the CNS, targeting α3 could lead to reduced side effects compared to other GlyR-α isoforms.

GlyR-α3 is produced as one of two possible splice variants, α3K or α3L. Post-transcriptional exclusion of exon 8 from the GlyR-α3 coding mRNA [3, 10–12] results in the α3K variant lacking 15 amino acids (TEAFALEKFYRFSDT) in the large intracellular loop between transmembrane α-helices TM3 and TM4 (Fig. 1A). GlyR-α3L is the predominant variant in a healthy brain, outweighing α3K approximately five-fold. Although both variants are always co-expressed in neurons, GlyR-α3K primarily localizes somatodendritically [5, 6, 13, 14], while α3L mostly localizes at the presynapse due to interaction with vesicular trafficking factor SEC8 [6], where it stimulates neurotransmitter release [6, 15–17], similar to other presynaptic chloride channels. Finally, neuronal cells additionally co-expressing GlyR-β endogenously will also contain postsynaptic heteropentameric α-β GlyRs, due to interaction with the postsynaptic scaffold protein gephyrin [18, 19].

**Fig. 1:**
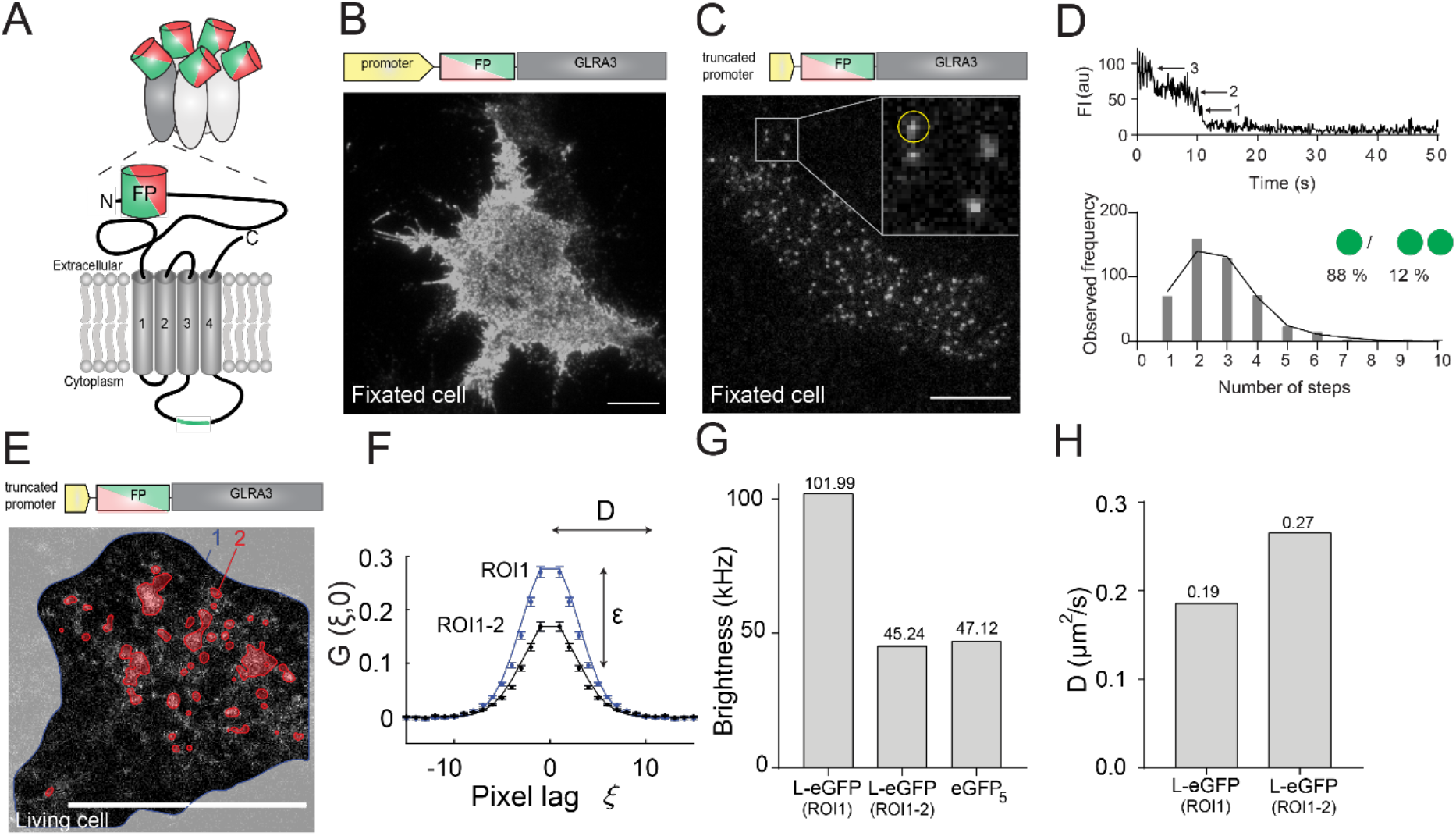
**Low copy number imaging of GlyR**-α3L-FP in HEK293 cells allows quantifying single pentamer properties. A) Subunit structure of GlyR-α3-FP with the fluorescent protein eGFP or mCherry (FP) fused to the terminus of the extracellular N-terminal domain and the position of the TM3-TM4 loop insert for GlyR-α3L in green. B) Representative cell with high copy number GlyR expression from a plasmid with a stong promoter. Scale bar, 10 μm. C) Representative fixated cell with low copy number GlyR expression using a plasmid with a truncated promoter. Scale bar, 10 μm. The inset is a magnification of the area indicated by the white square, white spots indicate single receptors, the yellow circle corresponds to the data in the upper part of panel D. D) Step-wise photobleaching subunit counting identify low numbers of fluorescent eGFP per fluorescent spot in the low copy number cells. E) Confocal fluorescence image of a live cell expressing GlyR used for RICS analysis. The edge of the cell is outlined in blue (region of interest, ROI1) and the high intensity clusters, automatically selected via frame-to-frame intensity thresholding (see Materials and Methods for more details), are highlighted in red (ROI2). Scale bar, 10 μm. F) The 1D section of the average 2D RICS autocorrelation function (the reader is referred to Fig. S3A-B for images of the 2D correlation functions) at spatial lag (ξ, 0) of a confocal image series of GlyR-α3L-eGFP expressing cells using either all pixels within ROI1 or within ROI1 minus ROI2. The reader is also referred to Video S1-2 for the different ROIs. The mean brightness *ε* and mean diffusion coefficient are determined from the amplitude and shape of the correlation function, respectively. G) Representative example of the molecular brightness (in kphotons/second) of diffusing GlyR-α3L-eGFP assemblies within ROI1 or within ROI1 minus ROI2, and, as a reference, molecular brightness of diffusing cytosolic eGFP_5_ measured as close to the bottom membrane as possible. H) Representative example of the diffusion coefficient of diffusing GlyR-α3L-eGFP within ROI1 or within ROI1 minus ROI2.

Previous reports have used fluorescence microscopy and electrophysiology to investigate the properties of homomeric GlyR-α3. Apart from their overall subcellular localization, fluorescence fluctuation imaging and single-particle tracking revealed that in live cells both (immunostained HA-tagged) splice variants exhibited free and confined diffusion in the membrane. Both fast (diffusion constant *D* ~0.1 μm^2^/s) and slow-diffusing (*D* ~0.01 μm^2^/s) species could be observed for both variants, with slow and confined diffusion being more prevalent for α3L than α3K [20, 21]. Fluorescence imaging using primary spinal cord or hippocampal neurons, or HEK293 cell lines, also evidenced that α3L is more prone to clustering in the cell membrane [5, 22, 23]. This suggests a role for the insert in the intracellular loop in directly promoting pentamer-pentamer interactions, whether or not combined with linking to immobile submembranous components that enhance the clustering process. It has also been shown using cell culture based whole-cell patch clamp experiments that, overall, α3L expressing cells exhibit slower desensitization kinetics than α3K [3, 24]. The longer TM3-TM4 intracellular loop seems to confer a larger stability to the α3L variant leading to slower desensitization, an effect that clustering seems to undo [25]. Finally, main-state single-channel conductances of 63-105 pS were observed for the α3L variant by different groups [25–29]. For the α3K variant, one group reported a similar conductance of 69 pS, suggesting the TM3-TM4 loop does not contribute to regulating the ion flux through the open channel [25].

Besides the molecular and functional differences of the α3 homomers described above, it is intriguing to know whether the two α3 splice variants can also form heteropentamers, and if they do, which effects this would have on GlyR function. Indeed, the pathological effect of the increased K-to-L expression ratio in TLE patients with a severe disease course hints to a functional direct interaction between the splice variants [5]. The existence of α3L/K heteropentamers in HEK293 cells was already suggested [22] via co-localization analysis of the differently labeled splice variants, although in this report the distinction between clusters of overlapping homopentamers or actual heteropentamers could not be made.

In this paper, we hypothesize that GlyR-α3 K/L splice variants functionally interact in a cell-biological context. If so, we would like to know what the molecular and functional properties of such heteropentamers are. We first develop strategies for expressing and imaging single GlyR-α3 pentamers in the membrane of live cells. We then use a combination of different quantitative fluorescence microscopy imaging and analysis methods including Pearson’s co-localization, raster/temporal image correlation spectroscopy [30, 31] and subunit counting via stepwise photobleaching [32, 33] to investigate the mobility, heteropentamerization and heterostoichiometry of co-expressed GlyR-α3 K/L splice variants. Then, we subject GlyR-α3 expressing cells to functional analysis via single-channel patch clamp. In the discussion section we integrate the results from the different types of experiments we performed and compare this with the present state of knowledge to better understand the cell-biological consequences of GlyR splice variant heteropentamerization.

## Results

### Advanced methodology for imaging single-pentamer properties of glycine receptors

Physiologically, GlyR-α3 molecules are present both as single pentamers and clusters of pentamers. As we were specifically interested in single pentamers, we first developed a cell-based fluorescent GlyR expression system and an analysis methodology that allows specifically analyzing the molecular properties of single GlyR-α3 pentamers in a way that is unbiased by clusters. First, to visualize the α3L and α3K isoforms of GlyR, we used plasmids encoding the GlyR N-terminally tagged with a green (eGFP) or red (mCherry) fluorescent protein (FP) (Fig. 1A, Fig. S1E-G) and transiently transfected these in HEK293 cells. These do not express GlyR endogenously but are known to be a relevant model system for studying GlyRs [3]. Immunocytochemistry (Fig. S1) and whole-cell patch-clamp electrophysiology (Fig. S2) confirmed a subcellular distribution and activity much like the endogenous situation, respectively, of the FP-tagged receptor.

Then, we followed a three-pronged approach to achieve the required low (single-molecule) and intermediate (10-100 nM) expression levels that are ideally suited for the planned single-pentamer analyses and for the diffusion analysis, respectively. We truncated the CMV promotor (similar to [34]), reduced the amount of GlyR encoding plasmid DNA while retaining the transfection efficiency via co-transfection with a non-coding plasmid [35] and limited the time between transfection and fixation or live-cell imaging. Using total internal reflection fluorescence microscopy (TIRFM) we imaged fixated cells expressing the normal- or low-expressing GlyR-α3L-eGFP plasmids (Fig. 1B-C). Indeed, with the latter plasmids we could easily find cells that clearly exhibited individual diffraction limited fluorescent spots (Fig. 1C), presumably single pentamers.

We next set out to prove whether these spots corresponded to single GlyR pentamers by recording time-lapse fluorescence images of transiently transfected cells and counting the number of eGFP molecules per diffraction-limited spot using single-spot photobleaching step measurements (Fig. 1D, top). As can be seen from the frequency distribution of the number of steps, a variety of bleaching steps ranging from 1 to 10 was observed (Fig. 1D, bottom). This has been observed before in bleaching experiments of GlyR-α1 in HEK293 cells [32, 36] and is attributed to a mixture of incomplete maturation of the fluorescent proteins, prebleaching of the eGFP, and single pentamers that are overlapping at a spatial scale smaller than the optical resolution. To analyze the data quantitatively, we fitted the distribution to a binomial model (Eq. 1, see Materials and Methods). This analysis resulted in a probability of 47% for eGFP to be maturated and unbleached and in 88% of spots not overlapping with other spots. Both of these values are similar to previous experiments on GlyR-α1 in HEK293 cells [32, 36]. This experiment thus suggests that in the low-expressing HEK293 cells, about 88% of detected fluorescent spots were likely single pentamers.

Finally, to corroborate that the majority of GlyRs detected in the cell membrane were indeed single pentamers, we used confocal raster image correlation spectroscopy (RICS). Practically, we performed experiments in cells with intermediate expression levels of GlyR-α3L-eGFP (ideally 10-100 nM [37]) (Fig. 1E). In such cells, we observed both regions with diffuse fluorescence, as well as regions with high-intensity fluorescent clusters, the latter presumably being GlyR aggregates that have been observed before [20]. After spatial autocorrelation of the images (Eq. 5) and fitting the resulting data to Eq. 6, we obtained both the molecular brightness ε (Eq. 7) and the mobility (diffusion constant, *D*) of the GlyR complexes diffusing in the membrane (Fig. 1F). The ε informs on the average number of fluorescing eGFP moieties in the diffusing complexes and, via comparison with a control protein, can be used to assess their average stoichiometry. The *D*, on the other hand, reports on the average size of these diffusing complexes, with slower diffusion indicative of larger complexes. When we included all pixels belonging to the cell membrane into the analysis (ROI1 in Fig. 1E, Fig. S3A and Video S1), the ε that we measured was significantly higher than that of a control protein, a cytosolic tandem eGFP pentamer (eGFP5) that we measured as close to the cell membrane as possible (Fig. 1G). When we additionally excluded the regions with an intense fluorescence signal (ROI1 minus ROI2 in Fig. 1E, Fig. S3B and Video S2), the brightness ε of the diffusing GlyR complexes was indistinguishable from that of the eGFP5 control Fig. 1G. Additionally, this experiment seems to show that properties of single GlyR pentamers can be specifically studied, in the case of intermediate-expression cells, by masking out regions containing clusters. The observed diffusion constant also depended on the ROI that was selected for the RICS analysis. Indeed, diffusion analysis in ‘ROI1 minus ROI2’ resulted in overall increased mobility, which directly proves the masking procedure efficiently removed the high-stoichiometry GlyR clusters (Fig. 1H).

In summary, we generated HEK293 cells expressing low amounts of GlyR-α3 splice variants labeled with fluorescent proteins and validated single-molecule and fluctuation imaging tools that allow focusing on the properties of single pentameric complexes excluding GlyR clusters. In the rest of the paper all analyses are performed on single GlyR pentamers, unless explicitly stated otherwise. Specifically, we take a closer look at the two splice variants, and at what happens when they are co-expressed in cells.

### Single homopentameric K and L variants exhibit a different diffusion signature

As a follow-up of the work of Notelaers et al. [20, 21], we next investigated the mobility of the two different splice variants GlyR-α3L-eGFP and GlyR-α3K-eGFP with RICS [38, 39] and temporal image correlation spectroscopy (TICS) [40], using image masking to specifically focus on single pentamers. RICS, which analyses μs-ms intensity fluctuations occurring within confocal image frames, is typically used to quantify the mobility of faster protein populations (*D* ≈ 0.1-500 μm^2^/s) while TICS, in which tens-of-milliseconds camera pixel intensity fluctuation are correlated over time, is typically used to quantify the mobility of proteins diffusing on a relatively slow timescale (*D* ≈ 0.001-0.1 μm^2^/s). Parallel application of both techniques allow identifying and characterizing different possible mobile protein populations [41]. Essential to this is choosing imaging conditions suited to the type of diffusion process (for RICS, see [42], for TICS, see [43]).

For RICS, we acquired confocal image series of living cells expressing either GlyR-α3L-eGFP or GlyR-α3K-eGFP at 37 °C as illustrated in Fig. 2A. Because in confocal microscopy the laser scans pixel per pixel and line per line while proteins diffuse, the resulting image will contain spatial fluorescence intensity fluctuations along any direction in the image, as depicted in Fig. 2B along the direction of a single line scan. We spatially correlated each image frame in the series (Eq. 5) and via fitting of the average spatial autocorrelation function (Fig. 2C-D, Eq. 6), we determined that the diffusion coefficients *D* of GlyR-α3L-eGFP (*D* = 0.26±0.11 μm^2^/s) and GlyR-α3K-eGFP (*D* = 0.29±0.08 μm^2^/s) were within experimental error the same (Fig. 2E). At least within the timescale of a single RICS image frame, the K and L variants thus exhibit similar diffusion.

**Fig. 2:**
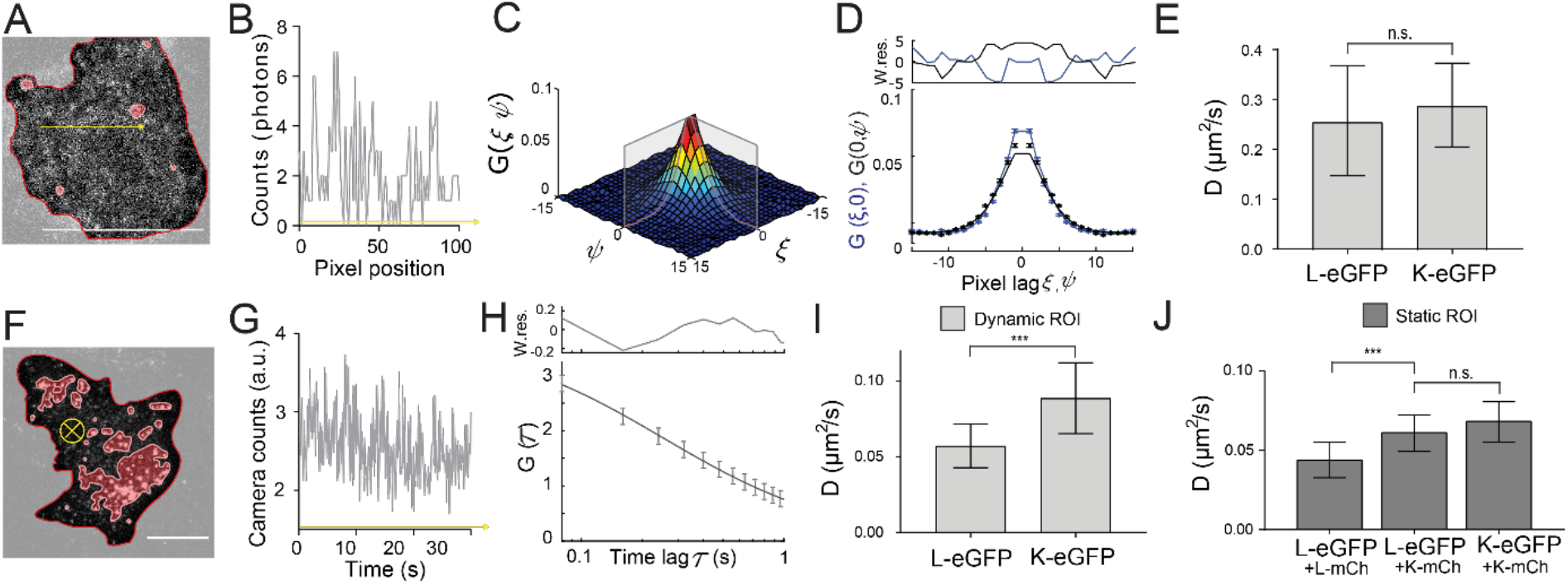
RICS and TICS evidence two diffusive subpopulations of single GlyR pentamers. A) Representative confocal microscopy image of the first frame from an image series of a HEK293 cell expressing GlyR-α3L-eGFP. Frame-based intensity thresholding was used to remove GlyR clusters and the extracellular region from the analysis. Scale bar 10 μm. B) Photon count values fluctuating along the yellow arrow in A. C) 3D autocorrelation with the grey outlining showing the average (ξ, 0) and (0, φ) autocorrelation function. D) Average (ξ, 0) and (0, φ) autocorrelation function and fit. Top graph displays the weighted residuals for the fit in the bottom graph. E) Average diffusion coefficient and standard deviation obtained via RICS for GlyR-α3L-eGFP and GlyR-α3K-eGFP. F) Representative TIRF microscopy image of the first frame from an image series of a HEK293 cell expressing GlyR-α3L-eGFP. Frame-based intensity thresholding was applied to remove GlyR clusters (indicated in red) and the extracellular region (indicated in light gray). Scale bar 10 μm. G) Camera count values fluctuating along the yellow arrow in F which is directed into the plane of the image to represent its direction through time. H) Average temporal autocorrelation function and fit. Top graph displays the weighted residuals for the fit in the bottom graph. I) The average diffusion coefficient and standard deviation obtained via TICS for single GlyR-α3L-eGFP and GlyR-α3K-eGFP. Here, a dynamic mask was used, calculated per frame, to omit both mobile and immobile GlyR clusters from the analysis. J) The average diffusion coefficient and standard deviation obtained via TICS for GlyR-α3L-eGFP when co-expressed with GlyR-α3K-mCh or GlyR-α3L-mCh compared to co-expression of GlyR-α3K-eGFP with GlyR-α3K-mCh. Image masking was based on the average intensity of the time series, so only static clusters were removed. Error bars on the bar graphs represent the standard deviation from *n* = 9-22 different cell measurements (Table S1-2). *** *p*-value < 0.001 obtained via an unpaired two sample *t*-test with unequal variance of the data.

For TICS, time-lapse images were acquired using TIRF-based widefield microscopy in living cells at room temperature (Fig. 2F). As the frame rate using a camera is much faster than for confocal microscopy, and oftentimes similar to the time it takes molecules to diffuse in and out of image pixels, fluorescence intensities tend to fluctuate from frame to frame due to molecular diffusion, as illustrated in Fig. 2G. By temporally autocorrelating each pixel’s fluorescence time trace (Eq. 10) and fitting a model to the obtained mean temporal autocorrelation function (Fig. 2H, Eq. 11), the diffusion coefficient can likewise be determined. In this way we obtained a diffusion coefficient of *D* = 0.089±0.023 μm^2^/s for GlyR-α3K-eGFP and a significantly lower diffusion coefficient of *D =* 0.057±0.014 μm^2^/s for GlyR-α3L-eGFP (Fig. 1I). First, this analysis reveals a second diffusive GlyR species, as values for *D* were significantly lower as observed with RICS, even when RICS measurement were performed at RT (Fig. S2F). More interestingly, however, the slower component of the L variant is significantly lower than the slow component of the K variant. To investigate the possibility that this could be related to inefficient removal of clusters from the analysis, which would affect the clustering-prone L variant more than the K variant, and thus also the observed mobility (Fig. S3C) [5, 22, 23], we performed a detailed comparison of different masking procedures (Fig. S3D). This showed a dependence of the observed *D* for both K and L on the type of mask used: whole cell (Video S3), static mask (Video S4, mask calculated on the average of all frames), dynamic mask (Video S5, calculated per frame), a significantly slower diffusion of the L variant was always observed. In other words, when looking at diffusing of single pentamers of GlyR-α3 on the slow TICS timescale, the L variant exhibits a slower mobility than the K variant.

Finally, we wanted to investigate whether co-expression of K would affect the mobility of L at the level of pentamers. Practically, we co-expressed GlyR-α3L-eGFP and the red mCherry FP-tagged version of the short GlyR isoform (GlyR-α3K-mCh) and performed single-color TICS on the acquired eGFP channel image series. Interestingly, we observed an increased diffusion coefficient for GlyR-α3L-eGFP in the presence of GlyR-α3K-mCh (Fig. 2J, Table S2, D = 0.061±0.01 μm2/s; the value is slightly different than in Fig. 2I because of the different mask used) as compared to cells co-expressing GlyR-α3L-eGFP and GlyR-α3L-mCh (Fig. 2J, Table S2, D = 0.044±0.01 μm2/s) or as compared to GlyR-α3L-eGFP alone (Fig. S3D, Table S2, D = 0.047±0.01 μm2/s). As expected, co-expression of GlyR-α3K-eGFP and GlyR-α3K-mCh (Table S2, D = 0.068±0.01 μm2/s) did not affect the mobility of the former as compared to GlyR-α3K-eGFP alone (Table S2, D = 0.074±0.01 μm2/s). These results are strongly indicative of a direct K-L interaction at the level of single pentamers, which we will investigate using dual-color imaging.

### Co-localization and co-diffusion proves GlyR-α3L/K heteropentamerization

To investigate whether GlyR heteropentamerization could be the cause of the increased mobility observed for GlyR-α3L-eGFP when co-expressed with GlyR-α3K-mCherry in the membrane of HEK293 cells, we recorded dual-color images via alternating-excitation TIRF microscopy (Fig. 3A).

**Fig. 3:**
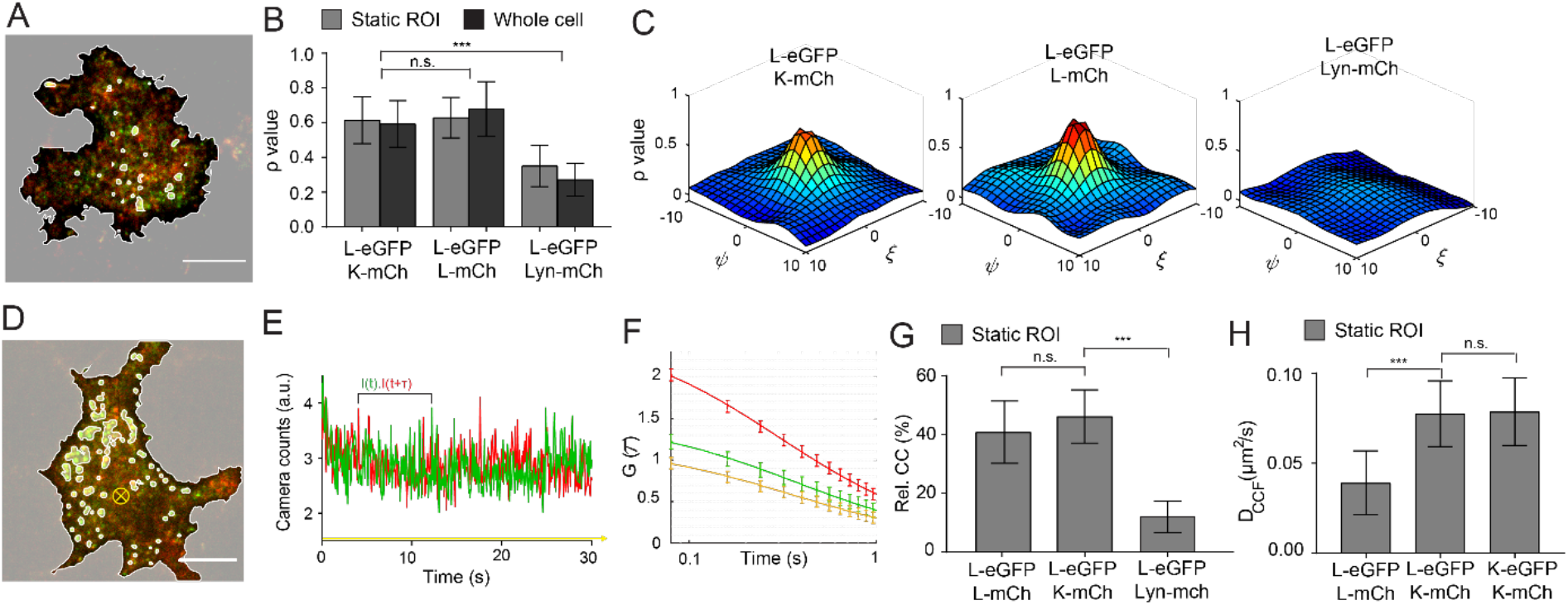
Co-localization and co-diffusion of GlyR-α3 isoforms in HEK293 cells at single-molecule expression confirms the presence of GlyR-α3L/K heteropentamers. A) Representative dual-color TIRF image of a HEK293 cell co-expressing GlyR-α3L-eGFP (green) and GlyR-α3K-mCherry (red). Using intensity thresholding over the average of the 5 first frames the cell membrane was selected and bright regions containing GlyR clusters were omitted. Scale bar 10 μm. B) Pearson’s correlation coefficient of cells expressing GlyR-α3L-eGFP and GlyR-α3K-mCherry and cells co-expressing GlyR-α3L-eGFP and GlyR-α3L-mCherry plasmids. Both experimental groups have significantly higher co-localization compared to the negative control with GlyR-α3L-eGFP and Lyn-mCherry. The *ρ* values are shown for cells including (grey) and excluding (dark grey) GlyR clusters. C) Spatial *ρ* from representative cells expressing GlyR-α3L-eGFP and GlyR-α3K-mCherry (left), GlyR-α3L-eGFP and GlyR-α3L-mCherry (middle) or GlyR-α3L-eGFP and Lyn-mCherry (right). D) Representative dual-color TIRF image of a HEK293 cell co-expressing GlyR-α3L-eGFP (green) and GlyR-α3K-mCherry (red). Intensity thresholding was applied over the average of the 400 frames (static ROI) to remove GlyR clusters and the extracellular region. Scale bar 10 μm. E) Dual-color fluorescence trace for one selected pixel over time (yellow arrow orthogonal to the image in D). F) Mean temporal autocorrelation (green and red) and cross-correlation (yellow) of all included pixels after intensity thresholding. Error bars are the 95% confidence intervals. G) The relative cross-correlation (Eq. 12) for cells expressing GlyR-α3L-eGFP and GlyR-α3K-mCherry, GlyR-α3L-eGFP and GlyR-α3L-mCherry plasmids, and the negative control with GlyR-α3L-eGFP and Lyn-mCherry. H) Average diffusion coefficient and standard deviation obtained via TICCS for cells co-expressing GlyR-α3L-eGFP and GlyR-α3K-mCh or GlyR-α3L-mCh, or for cells co-expressing GlyR-α3K-eGFP and GlyR-α3K-mCh. Error bars represent the standard deviation from *n* = 5-22 measurements (see Table S2-3 for *n*). *** *p*-value < 0.001 obtained via an unpaired two sample *t*-test with unequal variance of the data.

To quantify the similarity of the two images and hence the colocalization of the two splice variants in the membrane, we calculated the Pearson’s correlation coefficient *ρ* (Eq. 8). The *ρ* describes the degree of correlation between green and red channel pixel intensities of a dual-color image [44, 45]. The values of *ρ* can range from 1 to −1, with 1 a perfect correlation, 0 when there is no correlation and −1 for when there is an inverse relationship (exclusion) between the images. The pixels included in the analysis were confined to the region of the cell membrane since the extracellular region holds pixels with both low green and red intensity values which falsely increases the *ρ* value [46]. In addition, when cells contained regions with clustering GlyRs, these regions were also omitted by static ROI intensity thresholding to ensure Pearson’s analysis was performed only on the heteropentamer fraction. This revealed a positive Pearson’s correlation coefficient calculated for images of cells co-expressing GlyR-α3L and GlyR-α3K, similar to the one calculated for cells co-expressing GlyR-α3L labeled with eGFP and mCherry (Fig. 3B), and significantly higher than for the negative control cells co-expressing GlyR-α3L-eGFP and the monomeric membrane protein Lyn-mCherry that does not interact with GlyR. To confirm that the Pearson’s coefficient was indeed determined mainly by the fluorescent receptors, and less by cellular background, *ρ* was calculated as a function of the pixel shift between the images in the *x* and *y* direction (Fig. 3C). For cells expressing GlyR-α3L and GlyR-α3K a clear positive peak was seen, indicative of real co-localization. For cells expressing non-interacting GlyR-α3L-eGFP and Lyn-mCherry, this peak was generally absent or very small and wide (Fig. 3C, right), indicative of a specific co-localization.

Although Pearson’s correlation is an excellent qualitative proof for protein-protein co-localization, to more directly investigate the hetero-oligomerization of the slowly diffusing GlyR-α3 population we used dual-color cross-correlation TICS (TICCS) in HEK293 cells co-expressing GlyR-α3L-eGFP and GlyR-α3K-mCherry [31, 40]. For TICCS, image acquisition of the bottom membrane was performed using dual-color fast alternating TIRF-based excitation microscopy (Fig. 3D). For each pixel position in the image series, the fluorescence time traces (Fig. 3E) were temporally auto- and cross-correlated (Fig. 3F, Eq. 10). While the temporal autocorrelation and cross-correlation functions on their own allow determining molecular parameters such as mobility (Eq. 11), the relative cross-correlation additionally is a proof for their co-diffusion, and even a measure for the interaction affinity between them [47]. For cells co-expressing GlyR-α3L-eGFP and GlyR-α3K-mCherry we measured a high relative cross-correlation (Fig. 3G, Eq. 12) that was similar to cells co-expressing GlyR-α3L-eGFP and GlyR-α3L-mCherry. This is a result from a similar high interaction affinity. Note that even for constantly interacting or even covalently linked molecules the maximum interaction value is typically around 50-60% (it never reaches the theoretical 100%) since it is limited due to factors such as incomplete fluorescent protein maturation or the partial overlap between green and red microscope detection volumes [48]. As a negative control, we analyzed cells containing GlyR-α3L-eGFP and Lyn-mCherry (Fig. 3G). We observed a very low cross-correlation amplitude (Fig. S3E) and significantly lower relative cross-correlation.

In contrast to single-color fluctuation experiments, dual-color TICCS offers the additional possibility to focus specifically on the diffusion properties of the heteropentameric complexes containing both eGFP and mCherry fluorophores. In line with the single-color experiments, these data also show that GlyR-α3L/K complexes exhibit higher diffusion coefficients compared to GlyR-α3L-eGFP/mCherry homopentamers (Fig. 3H, Table S2). Interestingly though, the heteropentamer *D* derived from the cross-correlation function was completely indistinguishable from that of K homopentamers, indicating that the K splice variant dominates the diffusion pattern of K/L heteropentamers. Together, the Pearson’s correlation and dual color TICCS experiments prove that GlyR-α3L and GlyR-α3K are localizing and diffusing as a complex. As the masked analyses we perform allow focusing on GlyR pentamers, this must mean the GlyR-α3K and GlyR-α3L splice variants can heteropentamerize. Moreover, heteropentamer diffusion is dictated by the short-loop splice variant K. We next wondered whether these heteropentamers existed in a defined heterostoichiometry or not.

### The stoichiometry of GlyR heteropentamers depends on the relative subtype expression

To provide further insights into the stoichiometry of heteropentamers we performed two types of experiments: single-molecule step-wise photobleaching and molecular brightness analysis. For the first experiment, we performed continuous TIRFM single-molecule imaging of the eGFP labels in fixated cells co-expressing GlyR-α3L-eGFP and GlyR-α3K-mCherry and analyzed the resulting single-molecule traces with a step-finding algorithm to count the number of fluorescing eGFPs in a single complex (Fig. 4A). As the co-localization and fluctuation experiments showed that under such experimental conditions, these complexes are most likely heteropentamers containing both eGFP and mCherry fluorophores, it is expected that compared to samples containing GlyR-α3L-eGFP homopentamers (Fig. 1D), the number of eGFP moieties per complex should be lower. Indeed, the experimental data revealed a distribution with, on average, less eGFP subunits per spot compared to GlyR-α3L-eGFP homopentamers (Fig. 4B; Kolmogorov-Smirnov test p < 0.01).

**Fig. 4:**
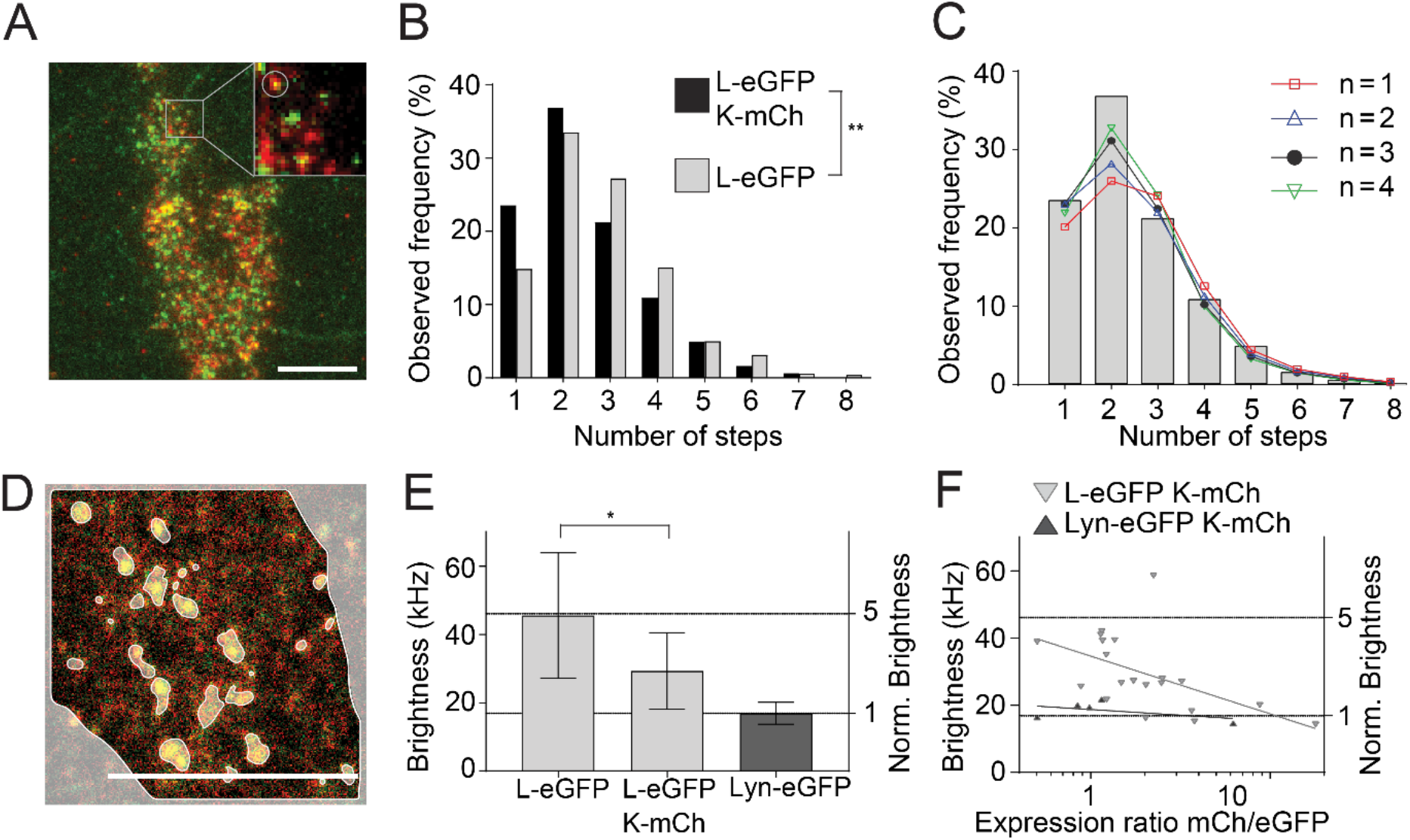
Automated subunit analysis and molecular brightness analysis shows the effect of relative expression on the GlyR stoichiometry. A) Representative images of HEK293 cells expressing GlyR-α3L-eGFP (green) and GlyR-α3K-mCherry (red). Scale bar 10 μm. B) Step distribution histogram of GlyR-α3L-eGFP in the presence (light grey, 301 spots) and absence (black, 477 spots) of GlyR-α3K-mCherry. In the presence of GlyR-α3K-mCherry there is a significant shift towards a lower number of GlyR-α3L-eGFP subunits. \*\**p*-value < 0.01 obtained with Kolmogorov-Smirnov test. C) Fitted binomial distribution functions with a sum of a 5^th^ order binomial and n^th^ order (*n*=1-4) binomial. See Table S4 for heteromeric fraction and *p*-value for the fit (χ2-test). D) Representative confocal microscopy image of the first frame from an image series of a HEK293 cell expressing GlyR-α3L-eGFP and GlyR-α3K-mCherry. Scale bar 10 μm. E) Molecular brightness comparison of the membrane-bound monomeric protein Lyn-eGFP to GlyR-α3L-eGFP in either the presence or absence of GlyR-α3K-mCherry. Error bars represent the standard deviation from *n* = 5-20 measurements (see Table S5 for *n*). * *p*-value < 0.05 obtained via an unpaired two sample *t*-test with unequal variance of the data. F) Molecular brightness of GlyR-α3L-eGFP (light grey) and monomeric Lyn-eGFP (dark grey) in the presence of variable amount of GlyR-α3K-mCherry. The semilog line fit shows a decrease in brightness for GlyR-α3L-eGFP upon increasing ratio of GlyR-α3K-mCherry to GlyR-α3L-eGFP.

We fitted the resulting step frequency distribution using two binomials, one representing the heteropentamer (up to 4 GlyR-α3L-eGFP subunits) and the other the homopentamer fraction (Eq. 2, Fig. 4C). The maturation probability (pm) and probability of overlapping spots (1-A) was fixed to 47% and 12%, respectively, based on the experiments on homopentamers (Fig. 1D), while the relative fraction of heteropentamers was fitted. The goodness-of-fit obtained via the χ2-test (Supplementary Table S4) was best for a 3rd order binomial and a heteropentameric fraction of 36% (goodness-of-fit *P*-value from a χ2 test = 0.725, with 1 being a perfect fit). However, as relatively good fits were obtained as well with a 2nd (*P*-value = 0.332) or 4th (*p*-value = 0.672) order binomial for a heteropentameric fraction of 23% and 67% respectively, these analyses are compatible with a scenario where heteropentamers contain on average 2-4 eGFP-containing subunits.

For the molecular brightness analysis, we recorded a confocal image series of living cells expressing GlyR-α3L-eGFP alone, or together with GlyR-α3K-mCherry (Fig. 4D) and analyzed the molecular brightness in the eGFP detection channel via dynamic-ROI based RICS (Eq. 7). As expected, this revealed a significantly lower molecular brightness for L/K heteropentamers as compared to L homopentamers (Fig. 4E, Table S5). Interestingly, the molecular brightness calculated in the eGFP channel scaled with the signal (count rate) ratio of GlyR-α3K-mCh compared to GlyR-α3L-eGFP (Fig. 4F), and a few cells even exhibited a similar molecular brightness as observed for the monomeric control Lyn-eGFP, meaning the presence of heteropentameric GlyRs with only a single L-eGFP subunit. Taken together, these experiments suggest the stoichiometry of GlyR-α3 heteropentamers is not fixed but variable, and depends on the relative expression of the L and K subtypes.

### The α3K electrophysiological signature dominates in heteropentamers

At this stage, we revealed the existence of K/L heteropentamers and investigated their molecular organisation. Lastly, we wanted to investigate possible functional differences between heteropentamers and homopentamers. Indeed, this might help to understand the consequences of an aberrant L/K ratio as observed in TLE. Practically, we used on-cell single-channel patch-clamp electrophysiology as opposed to whole-cell measurements to avoid averaging out activities of different co-existing species. Moreover, to verify the GlyR expression levels and ensure patch clamp measurements were performed under identical conditions as the fluorescence experiments, the patch clamp setup was mounted directly onto the single-molecules fluorescence microscope used for the stepwise photobleaching and TICS/TICCS experiments. Cells were transfected with either GlyR-α3L-eGFP or GlyR-α3K-mCherry, or with both. Importantly, the transfection conditions were similar to those used in the TICCS measurements, where we showed a high prevalence of heteropentamers for cells transfected with both plasmids. We selected cells with proper expression levels using fluorescence microscopy (Fig. 5A) and then performed on-cell single-channel electrophysiology to determine different possible conductance states (Fig. 5B).

**Fig. 5.**
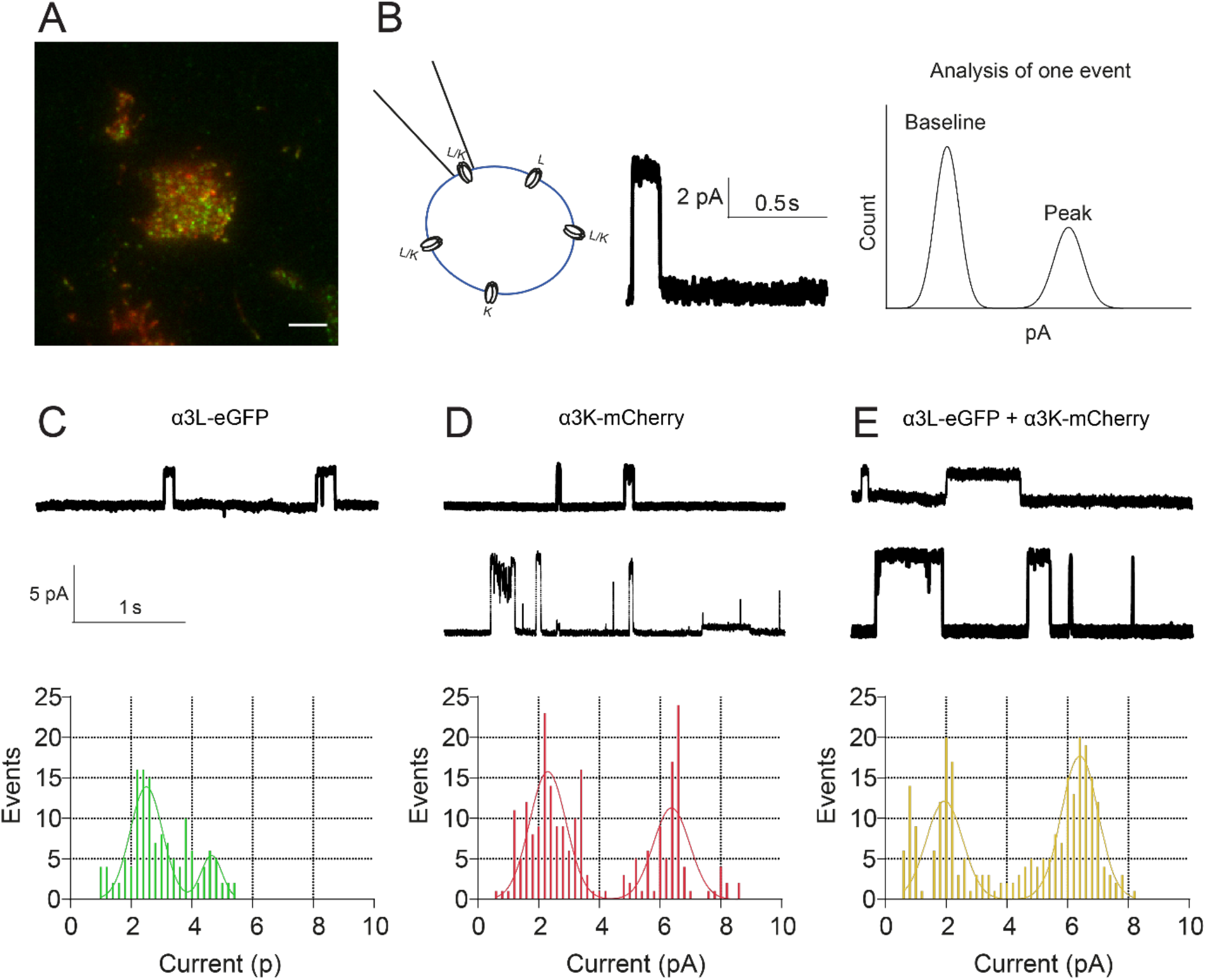
On-cell measurements show activated **α3K yields a high-current conducting state**. A) Transfected cells are identified by eGFP or mCherry fluorescence. Scale bar 5 μm. B) Example of a current time trace and analysis of a single opening event during which a peak current is seen. On-cell measurements are performed by placing a patch pipette to measure chloride currents through a single glycine receptor. For each single channel event, a histogram of all measured currents is fit with a double gaussian, yielding a best fit value for the mean baseline current and the mean peak current. Subtracting the mean baseline current from the mean peak current yields the ‘mean current of the single opening event’ C-E) Examples of single-channel current traces with accompanying histograms of ‘mean currents of the single opening events’ of cells transfected with α3L (*n* = 10, red), α3K (*n* = 18, blue) or α3L and α3K (*n* = 15, black). Histograms were fit with a double Gaussian to identify the most common peak amplitudes of single events. For α3K and for α3L and α3K an example of both a low and a high current single-channel trace is shown.

Like this, we could reveal two frequently occurring states for all tested conditions (Fig. 5C-E), with the first conductance state (around 2.3 pA) similar for all conditions (K, L and K/L), but the second conductance state showing significantly lower currents for α3L (around 4.7 pA) compared to α3K and α3L + α3K (around 6.4 pA). Contrary to what is known from [25–29] literature, our data seem to indicate that the presence of the α3K subunit in the GlyR yields higher currents.

## Discussion

The glycine receptor is a ligand-gated chloride channel that plays a crucial role in the general physiology of the CNS. Its α3 isoform, in particular, is involved epilepsy and chronic pain [5, 9, 26]. Two splice variants of α3 exist (Fig. 1A). As homopentamers, the splice variants differ in subneuronal distribution, electrical conductance and desensitization, clustering tendency, interactions with subcellular components and diffusion properties [3, 5, 6, 20–22], yet it is not known whether L and K variants can heteropentamerize. Indeed, such a process may lead to new or intermediate properties, or properties biased more towards one variant or the other. In an attempt to provide a more detailed fundamental cell-biological understanding of the working of GlyR-α3, we thus investigated the hypothesis that the different splice variants of GlyR-α3 can assemble into functional heteropentamers. To prove this hypothesis via advanced fluorescence imaging we first had to set up a new quantitative image analysis methodology for studying single pentamers in a complex sample, cell membranes containing both single pentamers and clusters of the same protein. Then, using this methodology we revisited prior work on the diffusion properties of GlyR-α3 splice variants in HEK293 cells to unequivocally prove whether RNA splicing determines the membrane mobility of the protein. Only hereafter could we embark on proving the existence of GlyR-α3L/K heteropentamers, and on quantifying their molecular and functional properties.

### Tools for studying defined molecular species in the case of oligomerization/clustering

A first methodological aim was to set up the necessary experimental tools to quantify single GlyR-α3 pentamer properties in cells. This is particularly challenging because of the tendency of GlyR-α3 to form subcellular clusters [5, 22, 23] that would overshadow the analysis. In previous research done by Notelaers et al., GlyR-α3 properties were investigated using fluctuation spectroscopy, single-molecule and super-resolution fluorescence methods, yet it was not explicitly investigated which observed species were representative of single pentamers or clusters [20–22]. Indeed, although image correlation spectroscopy (ICS) methods are quite robust in quantifying concentrations, diffusion and stoichiometry for monodisperse samples [49], they perform particularly badly in the case of polydisperse ones containing clusters, aggregates or multimeric species [50–52]. Here, we exploited the molecular fluorescence brightness of fluorescent protein labeled GlyR splice variants to validate that our methodology does specifically allow studying single pentamer properties. On the one hand, we performed subunit counting via stepwise photobleaching [32, 33] experiments in low-expressing cells in which fluorescent homomeric GlyR-α3L-eGFP was present as clearly discernable fluorescent spots (Fig. 1C-D). This way we found that the number of fluorescence bleaching steps per spot was similar to the previously studied GlyR-α1 under non-clustering conditions in HEK293 cells [32, 36]. On the other hand, we used a more recent extension of classical ICS called arbitrary-region ICS (ARICS) [50], where image series are segmented based on the local pixel fluorescence intensity, to specifically quantify the average molecular fluorescence brightness for single GlyR pentamer complexes diffusing in the live cell membrane (Fig. 1E-H; Video S1-2). With this analysis, we could show that results for (the more clustering-prone) homomeric GlyR-α3L-eGFP expressing cells were in line with non-clustering stoichiometric control proteins. This finally proved that our cellular expression system, HEK293 cells expressing fluorescent protein labeled GlyR splice variants from a crippled CMV promotor, and single-molecule photobleaching and segmented ICS analyses, are adequate for studying single pentamer properties, even when a non-negligible clustering subpopulation is present. Apart from the investigations performed in the rest of our paper, the methodological toolbox presented here can be applied for examining protein interactions, oligomerization, mobility and stoichiometry of other oligomeric receptors or multimeric proteins. Also, relative to the original methodological publication on segmented ICS [50] (detailed protocol in [53]), we did extend our in-house developed software for robust segmented single- and dual-color raster and temporal ICS analysis. This software can be downloaded free-of-charge (see Materials and Methods), is fully documented (https://pam.readthedocs.io/en/latest/mia.html), can be operated via a convenient graphical user interface from Microsoft and Apple operating systems, accepts a variety of images/videos, and can export figures and videos directly in publication-format.

### Specific subcellular interactions of single GlyR-α3L pentamers decrease their membrane mobility

Physiologically, GlyR-α3 is present in cells both as clustered and single pentamers. While α3K is more randomly distributed over the cell membrane, the situation for α3L is balanced somewhat more in favor of clusters. The α3L interacts with submembranous components specifically enriched at the presynapse, where it can promote (at glutamatergic nerve termini) neurotransmitter release [6]. Functionally, clustering of α3L thus seems to be an efficient way to promote this local enrichment. Notelaers et al. previously reported that overall, the subcellular mobility of α3L was lower than of α3K [20, 21]. For systems undergoing Brownian diffusion, the mobility (more specifically, the translational diffusion constant) of a freely-diffusing entity scales inversely with its size (Einstein-Smoluchovski relation). For membrane proteins in particular, mobility scales with the radius of the transmembrane region [54] [54, 55]. For the specific case of GlyR-α3L, receptor clustering would increase the size of the diffusing complex, and this would reduce mobility. Likewise, however, strong interactions of GlyR-α3 with large or immobile submembranous components would also likely reduce its overall mobility. As a follow-up of the work of Notelaers et al., we investigated whether a difference in single-pentamer mobility between the L and K variants can also be detected using our experimental setup. To this extent, we performed both confocal and TIRF-based microscopy and ICS analysis of GlyR-α3 expressing HEK293 cells to study the diffusion properties of the two splice variants. We segmented the images before ICS analysis to exclude those pixel regions containing GlyR clusters. Via confocal raster ICS (RICS) analysis we observed a fast freely diffusing component for both isoforms with similar diffusion coefficients (*D*α3L = 0.26±0.11 μm^2^/s and *D*α3K = 0.29±0.08 μm^2^/s) (Fig. 2A-E, Table S1). The existence of this freely diffusing component, that has been described before [21], suggests that at least a fraction of the GlyR-α3 population does not interact with immobile cellular components, or that the mere limited affinity for the latter defines the presence of a significant unbound component. The presence of functional GlyRs with relatively high mobility is, however, not surprising. It could allow for a faster reconstitution of non-desensitized GlyR receptor pools, as has been shown previously for the AMPA receptor, another ligand-gated ion channel [56]. When we studied the diffusion of single GlyR-α3 pentamers using TIRF-based temporal ICS (TICS), we observed a second, much less mobile species for both splice variants, which, interestingly, was even less mobile for α3L as compared to α3K (*D*α3L = 0.057±0.014 μm^2^/s and *D*α3K = 0.089±0.023 μm^2^/s) (Fig. 2F-I, Table S1-S2). This observation, in term, strongly suggests that the 15-residue-longer loop of GlyR-α3L relative to GlyR-α3K does indeed stabilize interactions with cellular interaction partners such as proteins or lipids, independent of GlyR clustering [6]. For primary neuron hippocampal cells, vesicular transport protein SEC8 targets the GlyR-α3L to the presynapse, and in vivo, GlyR-α3L was indeed detected at presynaptic terminals of glutamatergic and GABAergic neurons [6]. An interesting follow-up study would be to use site-directed mutagenesis of the insert region to more closely study sequence motifs of GlyR-α3L binding partners that control axonal receptor trafficking and localization. Conversely, GlyR-α3K is mainly distributed somatodendritically, but is also expected to be present in axonal and presynaptic compartments as this splice variant lacks a subcellular targeting signal and hence diffuses throughout the neuronal plasma membrane. This notion is furthermore supported by a recent study showing that there is no GlyR-β protein expression in hippocampal neurons [57], which could target the GlyR-α/β heteropentamers to postsynaptic gephyrin-positive scaffolds [19, 58].

### GlyR-α3L and GlyR-α3K splice variants form heteropentamers of variable stoichiometry

Co-clusters of GlyR-α3 splice variants have already been reported [22]. The single-color TICS experiments we performed in the present study, however, provided a first hint towards a direct interaction between α3L and α3K splice variants in the form of heteropentamers, since co-expression of α3K increased the mobility of single α3L pentamers (*D*_α3L_ = 0.047 ± 0.012 μm^2^/s and *D*_α3L+K_ = 0.061 ± 0.011 μm^2^/s) (Table S2). Hetero-oligomers of different isoforms of GlyR-α/β [59] and of other ion channels such as the NMDA receptors have been described before, and also the biogenesis of GlyR-α3 would be compatible with it [60]. For GlyR-α3 they are of specific interest because of the differential subcellular localization of splice variants [6] and because of their different electrophysiological desensitization signatures [3].

To provide a more conclusive answer, we first demonstrated co-localization between GlyR-α3L and GlyR-α3K upon co-expression in HEK293 cells using a spatial version of Pearson’s colocalization analysis that is more robust against coincidental pixel co-localization (Fig. 3A-C, Fig. S3H, Table S3) [44, 61]. Subsequently, we used dual color temporal ICS (TICCS) to unequivocally demonstrate, for the first time, heteropentamerization of GlyR-α3L and GlyR-α3K (Fig. 3D-G, Table S2). Finally, we employed direct subunit counting via stepwise photobleaching to quantify that the average stoichiometry of heteropentamers is 2-4 α3L-eGFP-containing subunits (Fig. 4A-C, Table S4). A non-negligible homomeric fraction was also present in all datasets, which furthermore supports the absence of a defined heterostoichiometry. Direct subunit counting via stepwise photobleaching was previously used to show that α1 and β isoforms, genetically labeled with fluorescent proteins, heteropentamerize in a α3β_2_ stoichiometry [59]. Finally, we carried out molecular brightness analysis to reveal that the heterostoichiometry is indeed variable and depends on the expression ratio of both splice variants (Fig. 4D-F, Table S5). Putting all stoichiometry data together we did not provide any evidence for a highly specific α3L/K stoichiometry.

### GlyR-α3L/K heteropentamers have GlyR-α3K-like mobility and conductance characteristics

The presence of heteropentamers can have several implications for GlyR-α3 function. In this paper, we investigated the subcellular mobility and electrophysiological signature of heteropentamers. Single-color TICS provides a readout of mobility, and evidenced that co-expression of α3K increased the mobility of α3L (Fig. 2J, Table S2). Of course, in the case of a subcellular mixture of homo- and heteropentamers, such single-color measurements only provide an average view, which is why we next performed a mobility analysis of only those species containing both α3K and α3L via image cross-correlation analysis via dual-color temporal ICS (TICCS) (Fig. 3H, Table S2). From these experiments it became apparent that the diffusion signature of the K isoform is dominant for the mobility of the heteropentamers. This additionally suggests that the subcellular interactions of α3L that render its mobility slow are multivalent rather than monovalent. As many as 5 units of α3L thus seem to be needed to result in its homomeric mobility signature. This might mean that the affinity of subcellular interactions of α3L is rather low, and that an avidity effect leads to the observed reduced mobility of homomers. Finally, combined on-cell patch clamp and fluorescence microscopy allowed us to investigate the single channel current of GlyR-α3 in cells expressing both GlyR-α3L-eGFP and GlyR-α3K-mCherry. A GlyR-α3K-shifted conductance was observed for cells containing heteropentamers, which was larger in amplitude compared to currents from GlyR-α3L-eGFP homopentamers.

As GlyR-α3L adopts the mobility signature of GlyR-α3K in heteropentamers, in regions of the brain where co-expression of GlyR-α3L and GlyR-α3K occurs, this could mean that heteropentamerization influences GlyR renewal in the plasma membrane, and as a result GlyR functionality. Consequently, this further stresses the importance of well-regulated alternative splicing for GlyR-α3 signaling. As in healthy people there is an increased presence of GlyR-α3L compared to GlyR-α3K, a small increase in alternative splicing would have an effect on even more GlyR-α3L pentamers due to heteropentamerization. Due to heteropentamerization a higher fraction of GlyR-α3L containing pentamers will have a higher mobility, which could enable faster reconstitution of the non-desensitized GlyR receptor pool [56]. The results from electrophysiology in particular also point to the possibility that the neuronal output can be increased by GlyR-α3 heteropentamers, particularly in conditions such as TLE where increased RNA editing and resulting gain-of-function receptors lever out rules of homeostatic regulation of the neuronal output [6, 14]. Importantly, subcellular trafficking and localization (pre- or postsynaptic, or e.g. in the distal and basolateral membrane compartments of epithelial cells) must be logically and interpretively distinguished from terms that describe single channel signatures of mobility and electrophysiology (conductance states). Indeed, a single receptor pentamer with specific mobility and conductance states can lead to very different outcomes depending on its subcellular localization. For example, due to its very small surface, the electrical capacity (C) of a presynapse is much lower compared to the somatodendritic compartment, and hence, one single channel conductance of chloride ions (Q) through the presynaptic plasma membrane will have a much greater impact on membrane potential (U) compared to the same conductance in the somatodendritic compartment (ΔU = Q/C).

## Conclusion

In this work, we investigated the long (L) and short (K) intracellular loop splice variants of the GlyR-α3 isoform, that is related to chronic pain and temporal lobe epilepsy. We unambiguously showed that these splice variants co-assemble into electrophysiologically active heteropentamers in live HEK293 cells. To do this, we had to set up and validate a combination of advanced single-molecule fluorescence, fluorescence fluctuation correlation and patch clamp methods, as the GlyR-α3 tends to cluster inside cell membrane, and this clustering is extraordinarily challenging for quantitative investigations. First and foremost, this work constitutes a methodological framework that can be used for investigating other types of complex hetero-oligomerizing molecular systems in a cell-biological context. Biologically, it turned out that, while the GlyR-α3L was well-known to determine the subcellular localization of GlyR-α3 channels, GlyR-α3K is leading in the regulation of both the in-membrane mobility of GlyR-α3, as well as in the ion channel’s activity. Indeed, heteropentamers were both more mobile than L homomers, and exhibited a larger open-state electrical conductance. Future research could be aimed at studying GlyR heteropentamer clustering, localisation and acitivity in primary neuron cells, as this would corroborate the importance of heteropentamers in neuronal signaling. Likewise, measuring channel open times would prove that heteropentamerization is important for fine-tuning of neuronal activity, which would, in turn, provide insights into the desensitization behaviour of heteropentamers.

## Materials and methods

### DNA plasmids

Plasmids encoding mouse GlyR-α3L or α3K containing an N-terminal eGFP or mCherry were already described [62] or obtained accordingly using standard molecular cloning technology by replacing mCherry with eGFP. N-terminal eGFP insert was amplified with PCR (5’-CGGTCTCCGGAATGGTGAGCAAGGGC-3’ and 5’-GGCCTCCGGACTTGTACAGCTCGTCCATGC-3’), the GlyR-α3L/K plasmids and the amplified eGFP insert were digested with BspE1. The vector plasmids were treated with calf intestine phosphatase before the ligation was performed. The enhancer region of the cytomegalovirus promotor in the GlyR-α3-coding plasmids was shortened similar as in [34] to reduce expression levels by mutagenesis. We did this by amplification of the GlyR-FP plasmids using PCR with primers 5’-ATATGGTACCTGGGAGGTCTATATAAGCAGAG-3’ and 5’ATAAGGTACCCCAGGCGGGCCATTTACCGTA-3’ followed by digestion with KpnI (ThermoFisher Scientific, Merelbeke, Belgium) and ligation using instant sticky-end ligase Master mix (NEB, Bioké Leiden, Nederland). Plasmids used as a negative control (Lyn-FP) were first used in [63] as a negative control for membrane receptor dimerization and encode the tyrosine-protein kinase Lyn coupled to a fluorescent protein eGFP or mCherry. Plasmids expressing eGFP or an oligomeric chain of 3 or 5 eGFPs (eGFP, eGFP_3_ and eGFP_5_) previously used in [64] were used as an stoichiometric reference.

### Cell culture and transfection

Human embryonic kidney 293 cells (HEK293 cells, provided by Dr. R. Koninckx, Jessa Hospital, Hasselt, Belgium) were cultured up to a maximum passage number of 20, at 37 °C and under a humidified 5% CO_2_ atmosphere in complete DMEM medium (D6429, Sigma-Aldrich, Overijse, Belgium) supplemented with 10% FCS (Sigma-Aldrich). At least 24 h before transfection, 150,000 cells were plated in complete medium in a 35-mm diameter #1.5 (170 μm glass thickness) glass bottom dish (MatTek, Bratislava, Slovak Republic). Cells were transfected via calcium phosphate-DNA co-precipitation [65]. The phosphate-DNA mix contained 86 μL HEPES-buffered saline (HBS) (280 mM NaCl, 10 mM KCl, 15 mM D-glucose, 1.5 mM Na_2_HPO_4_.2H_2_O, 50 mM HEPES, pH 7.1,) and 2000 ng total plasmid DNA per dish including the 50-1000 ng FP-tagged encoding plasmids supplemented with an empty plasmid vector (pCAG-FALSE, Addgene plasmid #89689) depending on the aimed fluorescence level [35]. To this mix 5.1 μL 2.5 M CaCl_2_ was added, and after 10 min of incubation at room temperature (RT) the mix was added dropwise to the cells.

### Immunostaining

Fixated cells expressing human GLRA3 were permeabilized at RT with permeabilization buffer (40 mL PBS, 2 g sucrose, 400 μL 10% Triton X-100) during 5 min, after which the cells were washed twice with washing buffer (40 mL PBS + 40 μL Triton X-100 10%). Next, cells were incubated for 10 min with proteinase K (0.1%, Thermo Fisher Scientific, Merelbeke, Belgium) for antigen retrieval. After washing once again with washing buffer, a blocking buffer (40 mL PBS + 0.4 g BSA + 40 μL Triton X-100 10%) was added to the cells. Finally, cells were incubated for 1 h with the primary anti-GLRA2 (1/2000 in blocking buffer; ab97628 Abcam, Cambridge, England), washed 3 times with blocking buffer and incubated for 2 h with Alexa 647 anti-rabbit (1/250, A21247, Thermo Fisher Scientific). After washing the cells 3 times with PBS, the cells were stored at 4 °C for limited time.

### Total internal reflection and widefield fluorescence microscopy imaging

A Zeiss ELYRA PS.1 inverted microscope with a Plan-Apochromat 100x/1.46 oil DIC M27 objective lens and Andor iXon+ 897 EMCCD camera operated at EM gain ~200 was used in total internal reflection fluorescence (TIRF) mode to selectively excite molecules near (< 200 nm) the bottom cell membrane. Images were recorded at room temperature using a multiband emission filter LBF 488/561 at a resolution of 256×256 pixels^2^ and a pixel size of 150 nm. The 488 nm and 561 nm HR diode-pumped solid-state lasers were used. Immunostaining imaging was done on the same setup, using an additional 642 nm HR diode-pumped solid-state laser. The reported laser powers were measured on the objective lens with immersion oil using a calibrated S170C microscope slide power sensor head (Thorlabs, Dortmund, Germany). Imaging was done using the ZEN software (Zeiss).

### Subunit counting by photobleaching analysis

TIRF images were acquired as decribed above using cells transfected with 50 ng GlyR-α3L-eGFP and 0-500 ng GlyR-α3K-mCherry which were fixated 22 h post-transfection for 24 h at 4 °C using 3% (w/V) paraformaldehyde in phosphate buffered saline. Before acquiring the images for the photobleachinig analysis, in each cell mCherry was photobleached with the 561 nm laser (5% power, 2.5 mW) in order to eliminate Förster resonance energy transfer between eGFP and mCherry. Next, 2000 frames were acquired at 100 ms per frame using the 488 nm laser at high enough power to induce step-wise photobleaching (1.5% power, 660 μW). Data analysis was performed using the Progressive Idealization and Filtering (PIF) software kindly provided by Dr. Rikard Blunck [32]. Molecules were located by selecting of 5×5 pixels^2^ spots with the signal-to-noise (δ*F/F*) setting at 20%. Next, intensity time traces were extracted from a 3×3 pixels^2^ region in the center of each spot. Partially overlapping spots were excluded from analysis. Photobleaching steps were identified via a step-finding algorithm when steps had a minimum length of 3 frames, and steps were not allowed to vary more than 60% in amplitude compared to other steps in the time trace. In addition, a minimal step signal-to-noise value of 2 was required. Cells with more than 10% accepted traces were included in the step frequency histogram. The step distribution of cells expressing only GlyR-α3L-eGFP was analyzed using the sum of two binomial distributions:

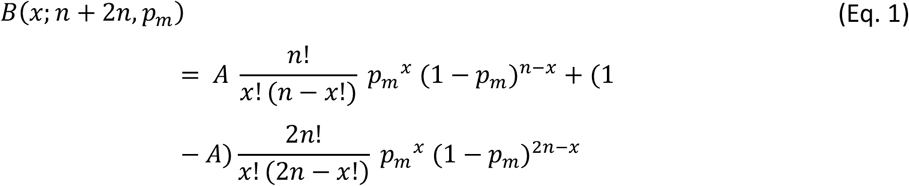

where *B* is the likelihood of observing *x* bleaching steps, *n* is the number of fluorescent eGFP molecules present in a single GlyR complex (*n* = 5), *p*_*m*_ is the probality that the fluorophore is maturated and non-bleached at the start of the recording, *A* is the fraction of spots containing not more than one GlyR complex and *1-A* is the fraction of spots containing two GlyR complexes. This equation assumes the fraction of spots containing more than two pentamers is negligible. In general, the *p_m_*-value reported in studies using subunit counting via stepwise photobleaching is typically on the low side (50-80%) [32, 33, 66] compared to other studies (~80%) [67, 68]. The broad range is appointed to variability between experimental groups such as the used cell line, fluorescent protein [69], temperature during maturation [32], cell fixation and fluorophore prebleaching [70].

To describe the step distribution of cells expressiong both GlyR-α3L-eGFP and GlyR-α3K-mCherry and determinte the stoichiometry (*het*) of the heteropentamers, Eq. 1 was extended to:

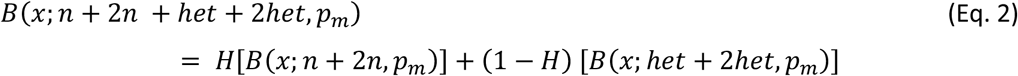

Here *H* represents the fraction of homopentamers and *1-H* represents the fraction of heteropentamers in the sample. Fitting this equation to the bleaching histograms of cells transfected with both, GlyR-α3L-eGFP and GlyR-α3K-mCherry, gives best fit values for *H* and *het*. Goodness-of-fit was determined using the chi-squared test. A good fit is indicated by a low χ^2^ value with p > 0.05, the model does not fit the data if p < 0.05 [59].

### Correlation analysis

Fluctuation imaging and co-localization analyses were performed in the software package PAM [71]. In all equations that follow, pre-processed intensity images *I*_*i*_(*x, y, t*) are converted into fluctuation images δ*I*_*i*_(*x, y, t*) prior to correlation analysis by subtracting the mean image intensity 〈*I*_*i*_〉:

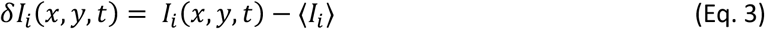

where *i* is the imaging channel, (*x, y, t*) denote the pixel coordinates and the angled brackets represent the average of all pixels included into the region-of-interest used for analysis.

### Raster image correlation spectroscopy

We used an inverted Zeiss LSM880 laser scanning microscope with a Plan-Apochromat 63x/1.4 Oil DIC M27 oil objective and MBS488/594 beam splittere to image live cells transfected with 100 ng GlyR-α3L-eGFP, 100 ng GlyR-α3K-eGFP or 50 ng Lyn-eGFP alone and/or combined with 0-1000 ng GlyR-α3K-mCherry, between 22-28 h post-transfection. Since RICS is ideally suited for capturing fast dynamics [31, 38, 41], the cells were held at 37 °C. However, to allow comparisons of RICS and TICS data, we did carry out limited RICS experiments at RT too (Fig. S3F). This revealed that the species observed with RICS still exhibited faster diffusion than those observed with TICS when measured at RT, and thus indeed represents a different subpopulation. Images were collected using parameters appropriate for RICS [42], i.e. 256×256 pixels^2^ with a 50 nm pixel size. Pixel dwell, line and image times were 8.19 μs, 4.92 ms and 1.26 s, respectively. The eGFP species were excited with a 488 nm argon-ion laser (0.3%, 1 μW) and mCherry species with a 594 nm HeNe laser (1%, 6 μW). Fluorescence was detected using a spectral detector (Zeiss Quasar) operated in photon counting mode in 23 spectral bins with ~9 nm bin width ranging from 490 nm to 695 nm. For quantitative analysis of eGFP-tagged molecules, bins 1-11 (490 nm to 589 nm) were summed for further analysis. Prior to autocorrelation analysis, we excluded contributions from slow processes such as cell and cell organelle movement using a moving average correction according to [39, 41, 72]:

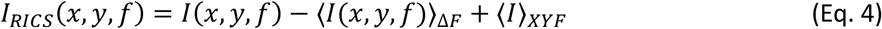

in which *I*(*x, y, f*) corresponds to each individual image, 〈*I*(*x, y, f*)〉_Δ*F*_ is the local mean image calculated over a short 3-frame interval from frame *f* − Δ*F* to frame *f* + Δ*F* with Δ*F* = 1, and 〈*I*〉_*XYF*_ is the mean intensity over all frames. Next, pixels outside the cell were removed by freehand-drawing based selection of the cell membrane, while GlyR clusters were removed using frame-based intensity thresholding. Specifically, both green and red images were first individually masked by intensity thresholding to remove (equalize to zero) pixels belonging to high-intensity clusters of fluorescence [50]. The final mask contained pixels that were included in each individual image’s mask and was smoothed using a 3×3 median filter as described above for co-localization analysis. Subsequently, the autocorrelation function was calculated per image frame using the arbitrary region-of-interest RICS (ARICS) algorithm [50]:

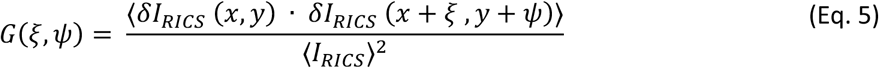

in which *ξ* and *Ψ* are the spatial lags in pixels, the · is the correlation operator, the angled brackets represent the average of all included pixels within the mask and 〈*I*_*RICS*_〉 is the average of all moving-average corrected pixels included into the region-of-interest used for analysis. To compare different datasets, we often plot only the (*ξ*, 0) correlations (example in Fig. 1F) or (*ξ*, 0) and (0, *Ψ*) correlations (example in Fig. 2D). Finally, the autocorrelation function was fitted with a one-component model assuming a two-dimensional Gaussian point spread function to obtain the diffusion coefficient, *D*, and average number of molecules in the focus, *N*.

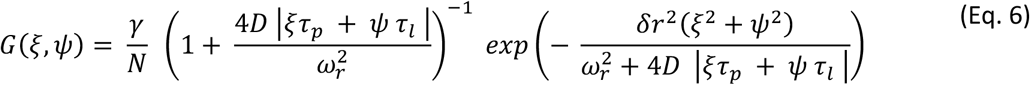

Here *γ* is the shape factor for a 2D Gaussian and equals 2^−3/2^ [73], *τ*_*p*_ and *τ*_*l*_ are pixel and line dwell times, δ_*r*_ is the pixel size and *ω*_*r*_ the lateral waist of the focus determined by calibration measurements (Fig. S4B). The RICS data was also used for calculating the molecular brightness of eGFP-containing diffusing molecules. Brightness (*ε*), expressed in kilophotons emitted per diffusing complex per second at the center of the confocal spot, was calculated by dividing the mean intensity of the image series (*F*) by the number of molecules obtained via RICS autocorrelation analysis (*N*_*ACF*1_)

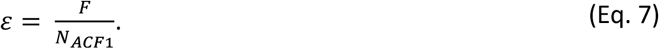

As stoichiometric references, cells were transfected with 5-10 ng eGFP, eGFP_3_ or eGFP_5_ encoding plasmids and investigated 22-28 h post-transfection as described above. When determining *N*, the moving average correction bias on the correlation amplitude was also corrected for as described before (Eq. 11 in [50]). Finally, stably focusing on the bottom membrane was achieved using a Zeiss Definite Focus.2 which acquired 60 frames at two different *z*-positions above the coverslip with an interval of 0.4 μm, alternating height each image frame, after which the time series at the *z*-position with highest average intensity was selected for analysis. We did also observe a clear effect of focus height above the coverslip on the molecular brightness, but not on the diffusion coefficient as shown in Fig. S3G, and as described before [41].

### Pearson’s co-localization analysis

A 400-frame TIRF image series of live cells transfected with 100 ng GlyR-α3-eGFP and 150 ng GlyR-α3-mCherry or 50 ng Lyn-mCherry was acquired 22h-28h post transfection at 80 ms per frame using alternating 2-color excitation. The eGFP species were excited during 20 ms at 488 nm (0.75% power, ~330 μW), followed by 20 ms excitation of the mCherry species at 561 nm (1.5% power, ~750 μW). A modified image correlation calculation was used to calculate the Pearson’s correlation coefficient *ρ* and to check the specificity of *ρ* [44, 61]. Image masking was performed as for RICS analysis. The *ρ* of the masked images was then calculated using:

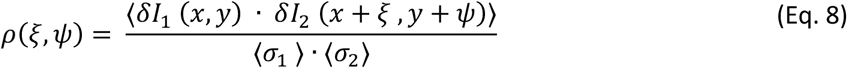

The *ρ*(0,0) is the classical Pearson’s coefficient, *ρ*≈1 means green- and red-labeled containing molecular complexes are overlapping, *ρ*≈0 means a random distribution and a value approaching −1 would mean exclusion. For the Pearson’s analysis, the same data as for fluctuation analysis was used, which contains significant shot noise. We therefore made an average of the first 5 image frames to obtain the most reliable Pearson’s correlation analysis (Fig. S3H).

### TICS and dual-color TICS

Sample preparation and TIRF image series recording was performed as described for the Pearson’s co-localization analysis. In each pixel the time series are preprocessed to remove the frame-to-frame variation of intensity using [41]:

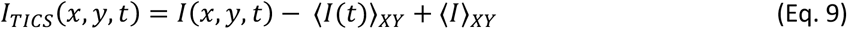

where *I*(*x, y, t*) is the intensity of any pixel, 〈*I*(*t*)〉_*XY*_ is the mean intensity of frame *t* and 〈*I*〉_*XY*_ is the mean intensity over all frames. The region inside the cell membrane was selected via freehand-drawing. To exclude high-intensity clusters either dynamic (as described above for RICS) or static image masking was applied (Videos S3-S5). For static region-of-interest (ROI) selection thresholding occurred based on the average intensity of the whole time series. Pixel-based auto- and cross-correlations were calculated using a one-dimensional discrete Fourier transform algorithm [40, 49]:

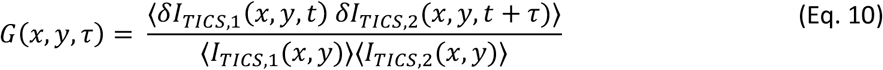

where *τ* is the time lag and δ*I*_*TICS*,1_ = δ*I*_*TICS*,2_ for autocorrelation of a single imaging channel, while for dual-color cross-correlation δ*I*_*TICS*,1_ and δ*I*_*TICS*,2_ are the values from the green and red image respectively. Finally, a one-component model for 2D diffusion was fitted to the autocorrelation functions (ACFs) and cross-correlation function (CCF) to obtain for each fit the average molecular diffusion coefficient, *D*:

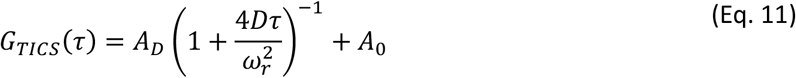

where *A*_*D*_ is the amplitude of the decaying part of the correlation function, *ω*_*r*_ is the radial waist of the point spread function (PSF) inherent to the resolution of the used microscope (Fig. S4A) and *A*_0_ is the offset caused by e.g. immobile molecules. To avoid influence of very slow motion (e.g. cell drift), the data was fitted until a 12-frame lag (i.e., ~1 s). The relative cross-correlation was obtained by dividing the amplitude of the cross-correlation function at the center by the mean of the two amplitudes of the corresponding autocorrelation functions.

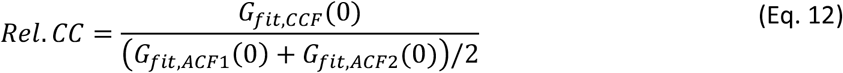

### Whole-cell patch-clamp electrophysiology

Cells were transfected with either GlyR-α3L-eGFP, GlyR-α3K-eGFP or GlyR-α3K-mCherry. Recordings were performed at room temperature in voltage-clamp mode using a HEKA EPC10 amplifier (HEKA Electronics, Lambrecht, Germany) controlled by HEKA acquisition software. Patch pipettes (3-4 MΩ) were filled with internal solution containing 120 mM CsCl, 2 mM Na_2_ATP, 2 mM MgATP, 10 mM EGTA and 10 mM HEPES, adjusted to pH 7.2 with CsOH. The standard external solution (SES) had a composition of 150 mM NaCl, 5.4 mM KCl, 2 mM CaCl_2_, 1 mM MgCl_2_, 10 mM glucose and 10 mM HEPES. Glycinergic currents were recorded at a holding potential *V*_*H*_ = −60 *mV*. Different glycine concentrations in SES including 20 μM, 50 μM, 100 μM, 200 μM, 500 μM were applied during 10 s. Maximum current amplitude was measured using FitMaster software (HEKA Electronics). The EC_50_ was calculated by plotting the normalized current as a function of concentration and fitting the data with the Hill equation (GraphPad Prism, La Jolla, CA, USA). For desensitization analysis, the decaying current phase was fitted using a mono-exponential in FitMaster software (HEKA Electronics, Lambrecht, Germany).

### On-cell single-channel electrophysiology

Cells were transfected with either GlyR-α3L-eGFP, GlyR-α3K-mCherry or both GlyR-α3L-eGFP and GlyR-α3K-mCherry. On-cell recordings were performed in voltage clamp mode at RT using a HEKA EPC10 amplifier. The external solution contained 120 mM NaCl, 4.7 mM KCl, 2 mM CaCl2, 1.2 mM MgCl_2_, 10 mM HEPES, 14 mM glucose, 20 mM TEA-Cl, 15 mM sucrose, adjusted to a pH of 7.4 with NaOH. Patch pipettes (5 – 15 MΩ) were filled with external solution and 30 – 80 μM glycine. The holding potential was set at +60 mV. Analysis of on-cell recordings was done using the FitMaster software. Amplitude histograms from single-channel openings were made by manually selecting single-channel opening events with a constant baseline. Histograms were fit with a gaussian fit yielding a mean open amplitude for the event. A histogram was made from all amplitudes in which the most frequently occurring conductance states were identified and fit with a gaussian.

### Summary of the Supplemental material

Fig. S1 shows the immunocytochemistry of the FP tagged GlyR. Fig. S2 illustrates the functional assessment of fluorescent protein tagged GlyR via electrophysiology, and shows the electrophysiology setup. Fig. S3 shows additional and control image correlation spectroscopy experiments. Fig. S4 shows focus size determination measurements of the Zeiss Elyra PS.1 and LSM880 microscopes. The supplementary tables include diffusion coefficients of the GlyR with frame-based thresholding (Table S1) and with average intensity-based thresholding (Table S2). Table S3 gives Pearson’s correlation coefficients to determine co-localization of GlyR-α3L and GlyR-α3K. Parameters obtaining from the bleaching histograms fits are in Table S4. Brightness of eGFP tagged proteins can be found in Table S5.

## Acknowledgements

We are greatly indepted to Mrs. Petra Bex and Dr. Sam Duwé for expert assistance with electrophysiology and microscopy, respectively. Prof. Em. Marcel Ameloot is thanked for critically reviewing the manuscript. Prof. Gonzalo E. Yevenes and Prof. Gustavo Morage-Cid (Department of Physiology, Faculty of Biological Sciences, University of Concepción, Concepción, Chile) are thanked for constructive discussions on the patch clamp experiments. Supervised students Hanneke Schroyen, Mahnoor Arif, Sam Vanspauwen and Jolien Broekmans are acknowledged for assistance in conducting experiments.

