## Supporting Information for "Glycine receptor α3K governs mobility and conductance of L/K splice variant heteropentamers"

* These authors contributed equally.

^#^ Correspondence:

### Supplementary figures

### Supplementary tables

### Supplementary figures

#### Supplementary Figure S1 – Immunocytochemistry of FP tagged GlyR

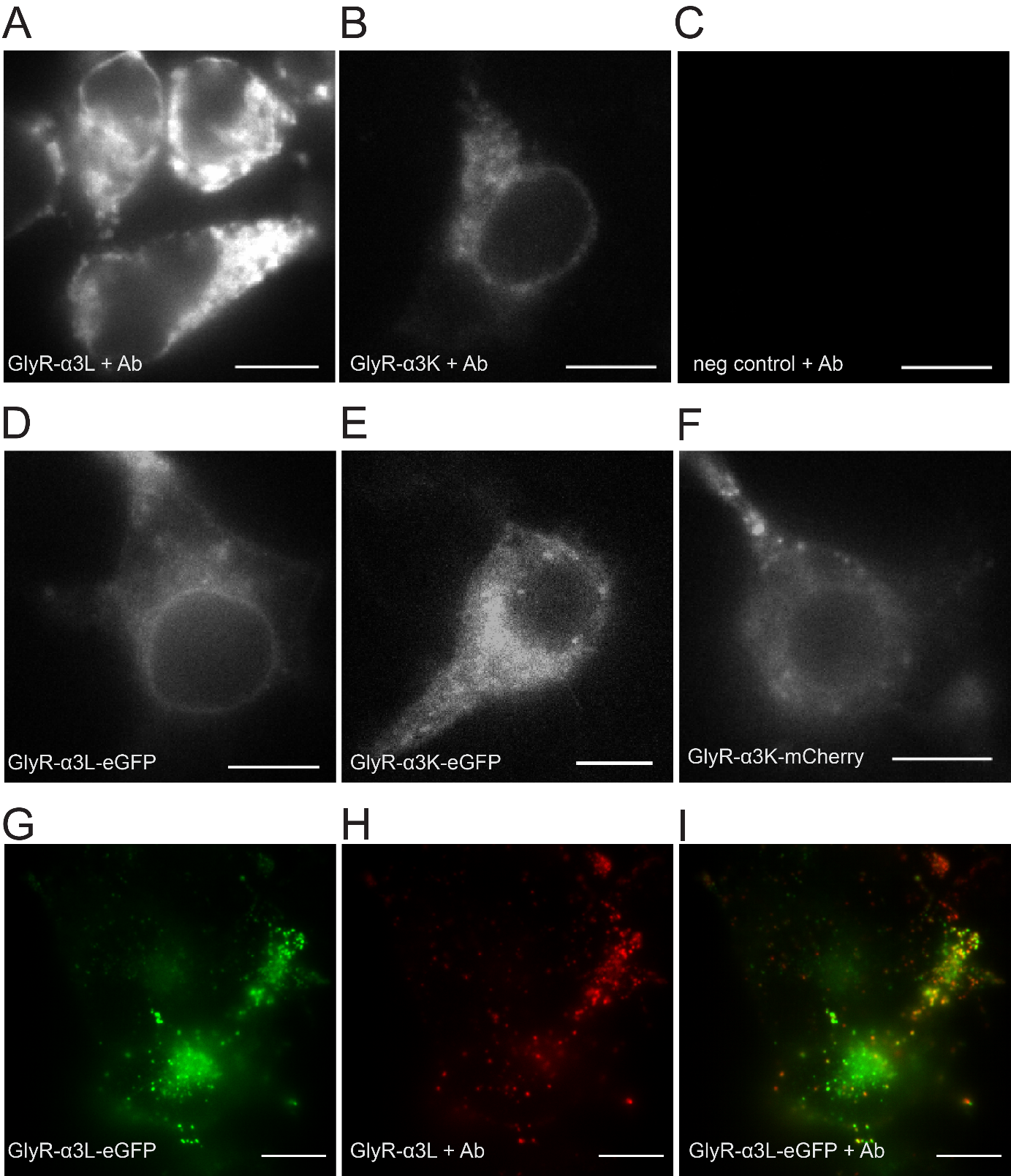

**Supplementary Fig. S1 –** **A-C)** Representative examples of immunocytochemistry of HEK293 cells expressing untagged (A) GlyR‑α3L and (B) GlyR‑α3K (middle), and (C) a negative control, HEK293 cells not expressing GlyR‑α, and stained with an anti-GlyRα antibody. Scale bars 10 µm. **D-F)** Representative examples of fluorescence imaging of HEK293 cells expressing (D) GlyR-α3L‑eGFP, (E) GlyR-α3K‑eGFP and (F) GlyR‑α3K‑mCherry. Scale bars 10 µm. **G-I)** Representative cell expressing the GlyR-α3L-eGFP construct (G), labeled with an anti-GlyRα antibody (see M&M) (H) and the overlap of both images (I).

#### Supplementary Figure S2 – Patch-clamp electrophysiology

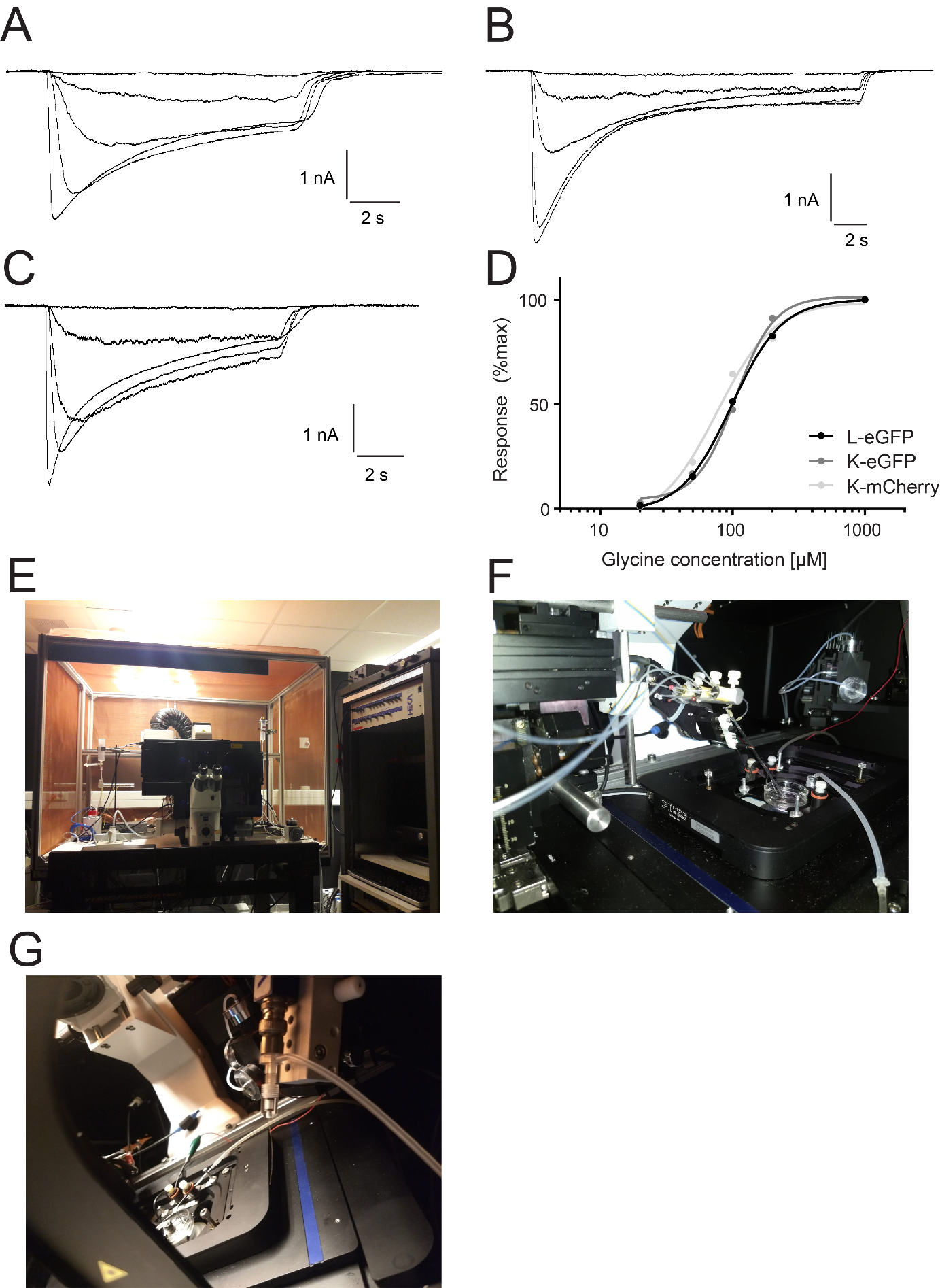

**Supplementary Fig. S2 –** **A-C)** Whole-cell patch‑clamp electrophysiology current responses of HEK293 cells transfected with plasmids encoding (A) GlyR-α3L‑eGFP, (B) GlyR-α3K‑eGFP and (C) GlyR-α3K‑mCherry. The responses were induced by 20, 50, 100, 200 and 1000 µM glycine. The glycine receptor currents show a pronounced dose‑dependent response and desensitization behavior. **D)** Half‑maximal responses (EC_50_) were obtained at glycine concentrations of 98.41 µM for GlyR-α3L‑eGFP, 104.8 µM for GlyR-α3K‑eGFP and 76.36 µM for GlyR‑α3K‑mCherry. These values are comparable to values reported in literature for mouse GlyR-α3 receptors expressed in HEK293 cells, for example an average EC50 value of 52 µM for GlyR-α3L and 47 µM for GlyR-α3K reported in (1). **E-G)** Patch-clamp electrophysiology setup (E) outside, (F) showing the fast-perfusion system that applies glycine solution to the cells for whole-cell patch-clamp, (G) showing the electrode connected to the patch-clamp amplifier.

#### Supplementary Figure S3 – Control experiments for image correlation spectroscopy

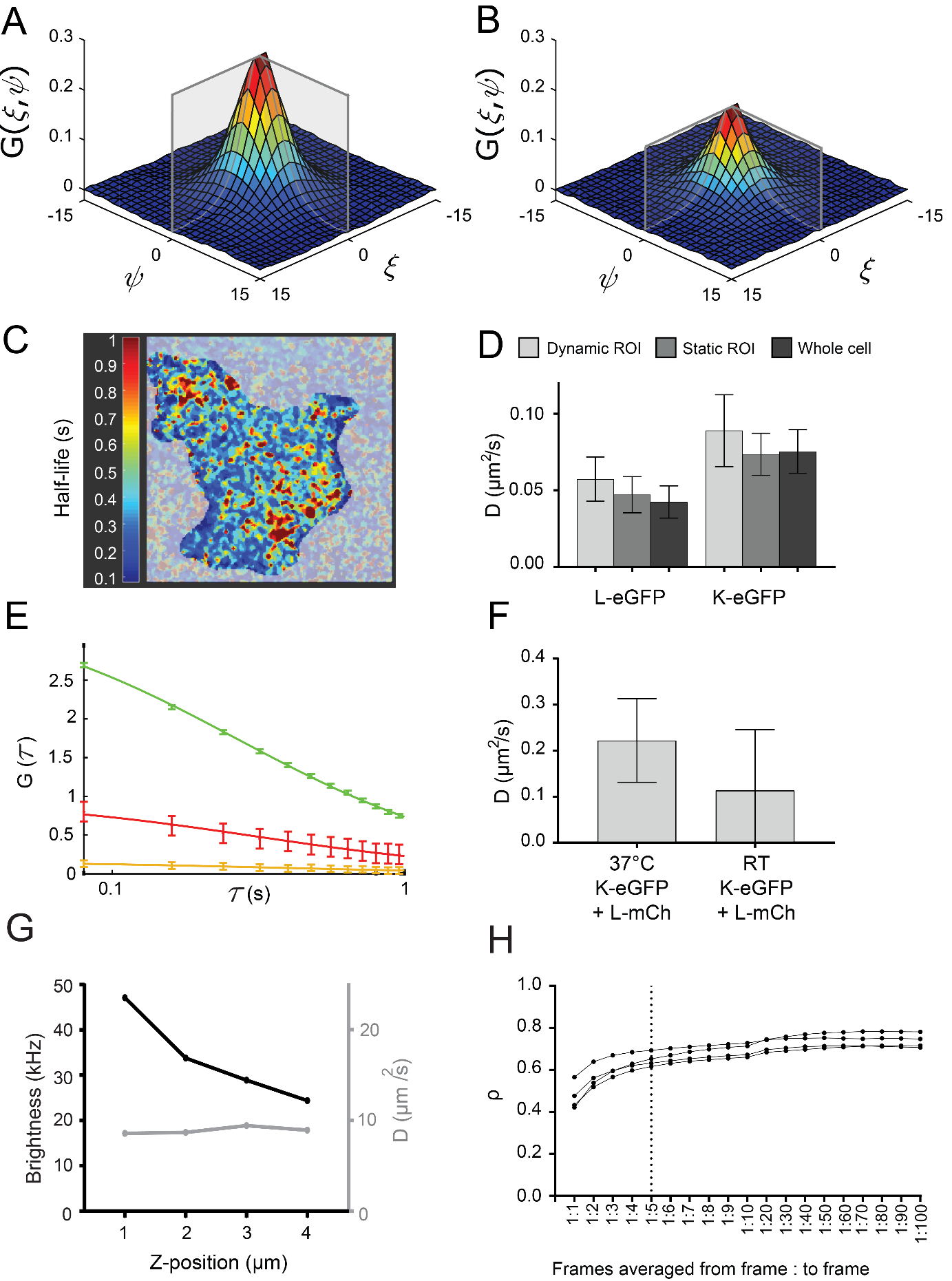

**Supplementary Fig. S3** **– A)** 2D correlation function of ROI1, supplementary to Fig. 1F. **B)** 2D correlation function of ROI1 minus ROI2, supplementary to Fig. 1F. **C)** Example of a spatially resolved TICS experiment. Shown is an image of the half-life of the TICS correlation function in each pixel, displaying regions with GlyR cluster exhibiting slow diffusion. **D)** Diffusion coefficient of GlyR-α3L eGFP and GlyR‑α3K‑eGFP obtained via TICS on the whole cell region (black), on the static ROI after excluding clusters with average intensity thresholding (dark grey) and on the dynamic ROI after excluding clusters with a mask dependent on the intensity in each single frame (light grey). **E)** Exemplary temporal mean autocorrelation (green and red) and cross‑correlation (yellow) of a cell expressing GlyR‑α3L‑eGFP and Lyn‑mCherry. Error bars are the 95% confidence intervals. Supplementary to Fig. 3. **F)** Temperature dependence of α3K-eGFP α3L-mCherry RICS autocorrelation data. **G)** Height dependence of RICS data of eGFP_5_, where 1 µm is axial position of maximum molecular brightness. **H)** The effect of averaging a variable number of frames for calculating the Pearson’s correlation coefficient *ρ* for 4 representative cells. When a single frame was used a low signal‑to‑noise in the image influences the *ρ*. When frames are averaged, the single‑to‑noise is increased resulting in a more constant *ρ*. We averaged 5 frames for calculation of the *ρ.*

#### Supplementary Figure S4 – Experimental optical resolution of the used microscopes

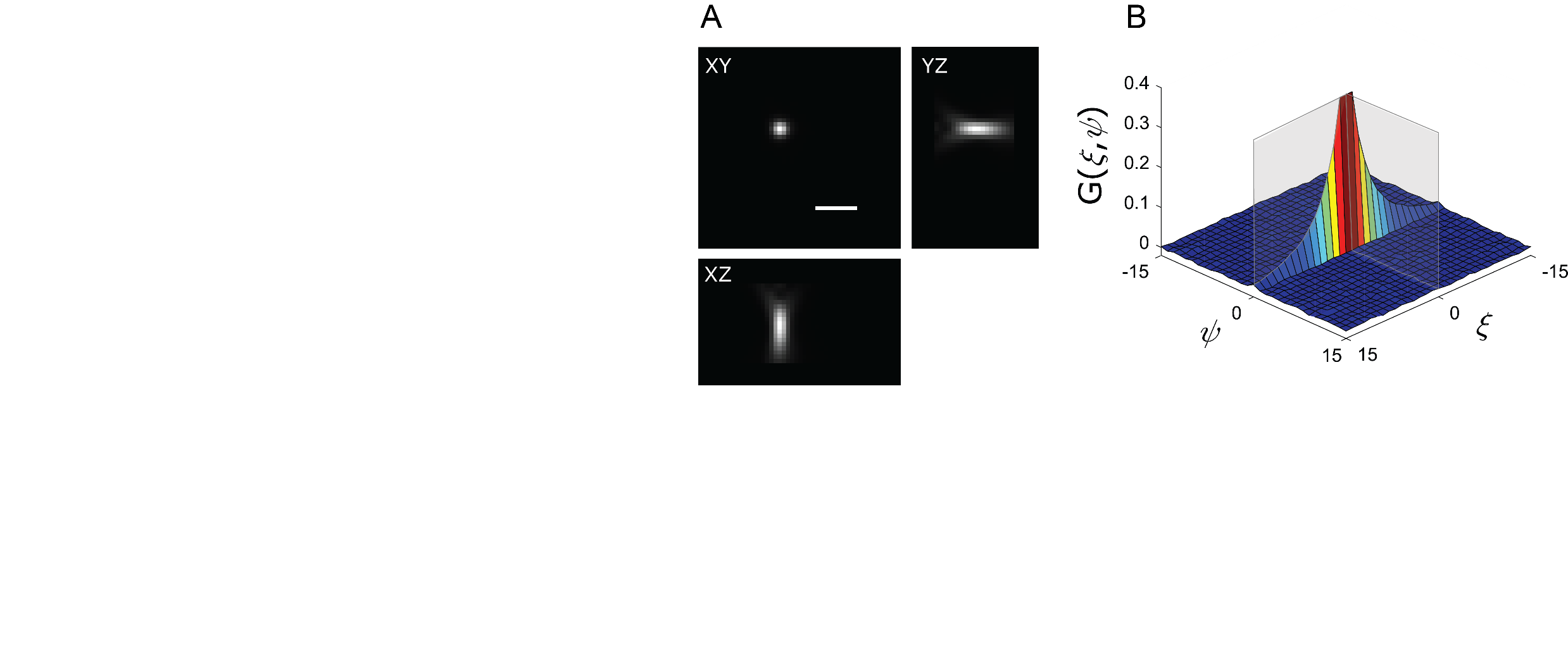

**Supplementary Fig. S4** – **A)** Experimental 3D point-spread function of the Elyra PS.1 microscope measured using 488-nm excitation on 20-nm fluorescent beads, used to calculate the lateral optical waist $\omega_{r}$ for TICS experiments. Scale bar, 1 µm. **B)** RICS autocorrelation function of ATTO488-COOH, used to determine the $\omega_{r}$ for RICS experiments (*D*_ATTO488-COOH,23°C_ = 373 µm^2^/s).

### Supplementary tables

#### Supplementary Table S1 – RICS and TICS dynamic ROI analyses

| **Method** | **Protein (eGFP)** | **n** | ***D ± SD* (µm^2^/s)** |
| --- | --- | --- | --- |
| **RICS** | GlyR‑α3L‑eGFP | 13 | 0.26 ± 0.11 |
| **RICS** | GlyR‑α3K‑eGFP | 9 | 0.29 ± 0.08 |
| **TICS** | GlyR‑α3L‑eGFP | 22 | 0.057 ± 0.014 |
| **TICS** | GlyR‑α3K‑eGFP | 13 | 0.089 ± 0.023 |

RICS and TICS of HEK293 cells expressing GlyR α3 eGFP isoforms with frame based thresholding (dynamic ROI). The diffusion coefficient (*D*) is calculated as described in Materials and methods. SD = standard deviation. The number of measured cells, that were not all measured on the same day, even within experimental groups, is indicated with *n*.

#### Supplementary Table S2 – TICS and TICCS static ROI analyses

| **Protein (eGFP)** | **Protein (mCherry)** | **n** | **D ± SD (µm^2^/s)** | |
| --- | --- | --- | --- | --- |
|  |  |  | **TICS (ACF1)** | **TICCS (CCF)** |
| GlyR‑α3L‑eGFP | - | 22 | 0.047 ± 0.012 | - |
| GlyR‑α3K‑eGFP | - | 13 | 0.074 ± 0.014 | - |
| GlyR‑α3L‑eGFP | GlyR‑α3L‑mCherry | 19 | 0.044 ± 0.011 | 0.039 ± 0.018 |
| GlyR‑α3L‑eGFP | GlyR‑α3K‑mCherry | 22 | 0.061 ± 0.011 | 0.078 ± 0.018 |
| GlyR‑α3K‑eGFP | GlyR‑α3K‑mCherry | 5 | 0.068 ± 0.013 | 0.079 ± 0.019 |

TICS and TICCS of HEK293 cells expressing GlyR α3 isoforms with average intensity based thresholding for clusters (static ROI). The diffusion coefficient (*D*) is calculated as described in Materials and methods. The number of measured cells, that were not all measured on the same day, even within experimental groups, is indicated with *n*.

#### Supplementary Table S3 – Pearson’s correlation coefficient

| **Protein (eGFP)** | **Protein (mCherry)** | **n** | ***ρ*** | |
| --- | --- | --- | --- | --- |
|  |  |  | **No clusters** | **Whole ROI** |
| GlyR‑α3L‑eGFP | GlyR‑α3K‑mCherry | 22 | 0.61 ± 0.14 | 0.59 ± 0.13 |
| GlyR‑α3L‑eGFP | GlyR‑α3L‑mCherry | 19 | 0.63 ± 0.12 | 0.68 ± 0.16 |
| GlyR‑α3L‑eGFP | Lyn‑mCherry | 11 | 0.35 ± 0.12 | 0.27 ± 0.09 |

The *ρ* is calculated as described in Materials and methods. The number of measured cells, that were not all measured on the same day, even within experimental groups, is indicated with *n*.

#### Supplementary Table S4 – Single-step photobleaching analyses

| **Heteromeric stoichiometry** | **Heteromeric fraction (%)** | **Homomeric fraction (%)** | **P‑value** |
| --- | --- | --- | --- |
| 1:4 | 0.13 | 0.87 | 0.024 |
| 2:3 | 0.23 | 0.77 | 0.332 |
| 3:2 | 0.36 | 0.64 | 0.725 |
| 4:1 | 0.67 | 0.33 | 0.672 |

Best fitted binomial distribution function consisting of a variable heteromeric fraction with nth order and a homomeric fraction of 5th order. P-values determined with the Chi^2^‑test.

#### Supplementary Table S5 – Brightness of eGFP-tagged proteins in HEK293 cells

| **Protein (eGFP)** | **Protein (mCherry)** | **n** | $\boldsymbol{\varepsilon}$ ***± SD* (kHz)** |
| --- | --- | --- | --- |
| GlyR‑α3L‑eGFP | - | 13 | 44 ± 19 |
| GlyR‑α3L‑eGFP | GlyR‑α3K‑mCherry | 20 | 29 ± 11 |
| Lyn‑eGFP | - | 11 | 17 ± 3 |
| Lyn‑eGFP | GlyR‑α3K‑mCherry | 5 | 18 ± 3 |

The brightness $\varepsilon$ is calculated as described in Materials and methods. The number of measured cells, that were not all measured on the same day, even within experimental groups, is indicated with *n*.
